# Biomass formation and sugar release efficiency of *Populus* modified by altered expression of a NAC transcription factor

**DOI:** 10.1101/2021.09.28.462238

**Authors:** Raja S Payyavula, Raghuram Badmi, Sara S Jawdy, Miguel Rodriguez, Lee Gunter, Robert W Sykes, Kimberly A. Winkeler, Cassandra M. Collins, William H. Rottmann, Jin-Gui Chen, Xiaohan Yang, Gerald A Tuskan, Udaya C Kalluri

**Author notes:** **Author for Correspondence:** Udaya C Kalluri. This manuscript has been authored by UT-Battelle, LLC under Contract No. DE-AC05-00OR22725 with the U.S. Department of Energy. The United States Government retains and the publisher, by accepting the article for publication, acknowledges that the United States Government retains a non-exclusive, paid-up, irrevocable, world-wide license to publish or reproduce the published form of this manuscript, or allow others to do so, for United States Government purposes. The Department of Energy will provide public access to these results of federally sponsored research in accordance with the DOE Public Access Plan(http://energy.gov/downloads/doe-public-access-plan).

## Abstract

Woody biomass is an important feedstock for biofuel production. Manipulation of wood properties that enable efficient conversion of biomass to biofuel reduces cost of biofuel production. Wood cell wall composition is regulated at several levels that involve expression of transcription factors such as wood-/secondary cell wall-associated NAC domains (WND or SND). In *Arabidopsis thaliana, SND1* regulates cell wall composition through activation of its down-stream targets such as MYBs. The functional aspects of *SND1* homologs in the woody *Populus* have been studied through transgenic manipulation. In this study, we investigated the role of *PdWND1B, Populus SND1* sequence ortholog, in wood formation using transgenic manipulation through over-expression or silencing under the control of a vascular-specific *4*-coumarate-*CoA* ligase (*4CL*) promoter. As compared to control plants, *PdWND1B*-RNAi plants were shorter in height, with significantly reduced stem diameter and dry biomass, whereas there were no significant differences in growth and productivity of *PdWND1B* over-expression plants. Conversely, *PdWND1B* over-expression lines showed a significant reduction in cellulose and increase in lignin content, whereas there was no significant impact on lignin content of down-regulated lines. Stem carbohydrate composition analysis revealed a decrease in glucose, mannose, arabinose, and galactose, but an increase in xylose in the over-expression lines. Transcriptome analysis revealed upregulation of several downstream transcription factors and secondary cell wall related structural genes in the *PdWND1B* over-expression lines that corresponded to significant phenotypic changes in cell wall chemistry observed in *PdWND1B* overexpression lines. Relative to the control, glucose release and ethanol production from stem biomass was significantly reduced in over-expression lines but appeared enhanced in the RNAi lines. Our results show that *PdWND1B* is an important factor determining biomass productivity, cell wall chemistry and its conversion to biofuels in *Populus*.

## Introduction

Woody biomass, harvested as feedstock for the pulp and paper, bioproduct and biofuel industries, is formed by tightly regulated biological and molecular genetic xylogenesis mechanisms. Primary xylem is formed from procambium while secondary xylem is formed from vascular cambium during secondary growth. The major constituents of secondary cell walls are cellulose, lignin, and hemicellulose (Darvill *et al*., 1980). Cellulose is the most abundant polymer in plants and is a polymer of glucose synthesized on the plasma membrane by the cellulose synthase (CesA) complex (Doblin *et al*., 2002). Lignin is the second most abundant polymer and is composed of guaiacyl (G), syringyl (S), and p-hydroxyphenyl (H) units derived through the phenylpropanoid pathway (Boerjan *et al*., 2003). In addition to cell division and expansion that occurs in primary cells, secondary xylem formation includes secondary wall deposition, lignification, and programmed cell death (Plomion *et al*., 2001). The formation of xylem cell walls is coordinately regulated at multiple layers by dozens of structural genes and transcription factors.

The major transcription factors that regulate secondary cell wall synthesis include SHINE (SHN), NAC (which stands for *NAM, ATAF1/2* and *CUC2*) domain transcription factors, and MYBs (Yamaguchi and Demura, 2010; Zhong *et al*., 2006). SHN is the master switch that controls the expression of down-stream transcription factors, NAC and MYBs (Ambavaram *et al*., 2011). Over-expression of *AtSHN* in rice (*Oryza sativa*) increased cellulose and decreased lignin (Ambavaram *et al*., 2011). The master and downstream transcription factors in the secondary cell wall transcription factor hierarchy include wood or secondary wall associated NAC domains (*WND/SND*), NAC secondary wall thickening promoting factor (NST), and vascular-related NAC domain (VND) transcription factors (Yamaguchi and Demura, 2010; Lin et al. 2017). The protein structure of NAC domain members is highly conserved in the N-terminal region and is required for nuclear localization and homo- or hetero-dimerization (Olsen *et al*., 2005). The C-terminal region has two conserved motifs, the LP-box and the WQ-box that regulate transcriptional activation (Ko *et al*., 2007; Yamaguchi *et al*., 2008). There is evidence for role of NAC family members in multiple plant processes and these functional roles can be redundant among sequence homologs (Aida *et al*., 1997; He *et al*., 2005; Hibara *et al*., 2003).

The NAC domain transcription factor is one of the largest families, with ~ 100 genes in *Arabidopsis* and soybean (*Glycine max*) and ~ 140 genes in rice (*Oryza sativa*) (Ooka *et al*., 2003; Pinheiro *et al*., 2009). In *Populus*, there are 163 genes clustered in 18 subfamilies (Hu *et al*., 2010). Among these, a few candidate transcription factors have been functionally characterized in model species such as *Arabidopsis* and rice. In *Arabidopsis*, at least three NAC domain members, *NST1, NST2*, and *NST3/SND1*, have been shown to have functional roles in regulating secondary cell wall biosynthesis (Mitsuda *et al*., 2007; Mitsuda and Ohme-Takagi, 2008; Zhong *et al*., 2006). T-DNA knockout mutants of *AtSND1* showed no difference from wildtype suggesting that the other isoforms might have compensated for the loss (Zhong *et al*., 2006). In contrast, either over-expression or dominant repression of *AtSND1* results in plants with weak stems and drastically reduced interfasicular fiber and xylary fiber wall thickness (Zhong *et al*., 2006). Over-expression of *AtSND1* resulted in massive deposition of lignified secondary cell walls suggesting that normal levels of *AtSND1* transcripts are necessary for maintaining proper cell wall thickening in secondary stems (Zhong *et al*., 2006). The defective secondary cell wall formation phenotype observed in *Arabidopsis snd1nst2* double mutants was restored by complementation with WNDs from *Populus*, suggesting that *Populus* WNDs regulate secondary wall biosynthesis (Mitsuda *et al*., 2007; Zhong *et al*., 2010; Zhong *et al*., 2007a). The NAC transcription factors bind to SNBE (secondary wall NAC binding elopements) in the promoters of its downstream targets and regulate their expression. *PtWND2B* induces expression of several wood associated MYB transcription factors and genes involved in secondary cell wall biosynthesis (Zhong *et al*., 2011; Mc Carthy et al. 2011). Over-expression of another NAC transcription factor gene, *Ptr-SND1-B1*, in *Populus* stem-differentiating xylem (SDX) protoplasts was reported to induce 178 differentially expressed genes (DEGs) of which 76 were identified to be its direct targets (Lin et al., 2013). Furthermore, two splice variants from NAC and VND transcription factor families are involved in reciprocal cross-regulation during wood formation (Lin et al., 2017). However, much less is known about the role of these transcription factors in maintaining cell wall composition. Recently, over-expression of a NAC family member, *PdWND3A*, was reported to affect lignin biosynthesis, decrease the rate of sugar release and reduce biomass (Yang et al., 2019). Given there is redundancy reported among functional roles of some NAC transcription factor family members and the knowledge of upstream master regulators of secondary wall biosynthesis, AtSND1, in *Arabidopsis*, here we sought to characterize the role of sequence ortholog, PdWND1B, in *Populus deltoides* in the context of biomass formation. To advance our knowledge on the role of additional NAC/WND transcription members in secondary cell wall biosynthesis, we developed transgenic *Populus deltoides* plants with xylem-specific over-expression or RNAi mediated silencing of *PdWND1B*, Potri.001G448400; *WND1B* has previously been referred to as PNAC017, VNS11, SND1-A2 (Ohtani et al. 2011; Zhong et al. 2010; Li et al. 2012; Hu et al. 2010; Takata et al. 2019). RNAi transgenic plants displayed weaker stems and altered cell wall composition as compared to control plants. Over-expression lines showed increased lignin content and significantly reduced ethanol production from stem biomass as compared to control plants. Our results confirm that *WND1B* plays an important role in secondary cell wall biosynthesis.

## Methods

### Phylogenetic analysis

Protein sequences of *Populus* WND isoforms were retrieved from *Phytozome v9.1: Populus trichocarpa v3.0* (Tuskan *et al*., 2006) and those corresponding to other plant species were obtained from NCBI. Phylogenetic analysis was performed in MEGA (Molecular Evolutionary Genetics Analysis) using the Neighbor-Joining method (Tamura *et al*., 2011). Bootstrap values were calculated from 500 independent bootstrap runs. Protein sequence alignment was performed using ClustalW and shading and percent similarity were predicted by GeneDoc (Nicholas *et al*., 1997).

### GFP localization

The full length coding regions of *PdWND1A* (Potri.011G153300) and *PdWND1B* (Potri.001G448400) were amplified from a *P. deltoides* xylem cDNA library (primers listed in Supplemental file 1) using Q5 High-Fidelity DNA polymerase (New England Biolabs, Ipswich, MA) and cloned in a pENTR vector (Invitrogen, Carlsbad, CA, USA). After sequence confirmation, the coding region fragment was recombined into a Gateway binary vector pGWB405 (Tsuyoshi *et al*., 2009) using LR clonase (Invitrogen). Plasmid from a positive clone was transformed to *Agrobacterium tumefaciens* strain GV3101. Tobacco infiltration and protein localization were performed as described previously (DePaoli *et al*., 2011; Sparkes *et al*., 2006). *Agrobacterium* harboring the binary constructs *PdWND1A* or *PdWND1B* were cultured overnight in LB media. After brief centrifugation, supernatant was removed and the pellet was dissolved in 10 mM MgCl_2_. The culture was infiltrated into four-week-old tobacco leaves. After 48 h, roughly 4 mm^2^ leaf sections were cut and fixed in 3.7% formaldehyde, 50 mM NaH_2_PO_4_, and 0.2% Triton X-100 for 30 min, rinsed with phosphate-buffered saline (PBS), and stained in DAPI (4,6′-diamidino-2-phenylindole, 1.5 μg ml^−1^ in PBS) for 30 min. GFP visualization and imaging was performed on a Zeiss LSM710 confocal laser scanning microscope (Carl Zeiss Microscopy, Thornwood, NY) equipped with a Plan-Apochromat 63x/1.40 oil immersion objective.

### Plant materials

The over-expression construct was developed by amplifying and ligating the 1235 bp coding region fragment of *PdWND1B* (gene model: Potri.001G448400, primers presented in Supplemental file) under the control of a vasculature specific *4-coumurate CoA-ligase* (*4CL*) promoter. The RNAi construct for the same gene was developed by amplifying a 300 bp coding region fragment (from 800 to 1100 bp) and ligated in sense and antisense orientation to form a hairpin with the chalcone synthase intron, under the control of *4CL* promoter. The binary constructs were transformed into wild-type *P. deltoides* ‘WV94’ using an *Agrobacterium* method (Caiping *et al*., 2004). Transgenic plants and empty vector transformed control plants that were roughly 10 cm tall were moved from tissue culture to small tubes with soil. After 2-months, plants were moved to bigger pots (6 liter) and were propagated in a greenhouse maintained at 25°C with a 16 h day length. At the time of harvest (six-month-old plants), plant height was measured from shoot tip to stem base, and diameter was measured two inches from the base of the stem. The bottom 10 cm stem portion was harvested, air-dried, and used for carbohydrate composition, cellulose, lignin, S:G ratio, and sugar and ethanol release analysis. Initial studies were performed on 10 transgenic lines for each construct and additional studies were performed on two to four selected lines. Data presented here is from two representative lines. Young leaf (leaf plastochron index, LPI-0 and 1), mature leaf (LPI-6^+^), and stem (internode portion between LPI 6 and 8) were collected, frozen in liquid nitrogen, and stored at −80°C until they were processed further.

### RNA extraction and gene expression studies

RNA from frozen ground stem samples was extracted using a Plant Total RNA extraction kit (Sigma, St Louis, MO) with modifications to the kit protocol. Briefly, 100 mg of frozen ground tissue was incubated at 65°C in 850 μl of a 2% CTAB + 1% βme buffer for 5 min, followed by the addition of 600 ul of chloroform:isoamylalcohol (24:1 v/v). The mixture was spun at full speed in a centrifuge for 8 min after which the supernatant in the top layer was carefully removed and passed through a filtration column included in the kit. The filtered elutant was diluted with 500 μl of 100% EtOH and passed through a binding column. This was repeated until all of the filtered elutant/EtOH mixture was passed through the binding column. Further steps including on-column DNase digestion (DNase70, Sigma), filter washes, and total RNA elution were followed as per the manufacturer’s protocol. cDNA was synthesized from 1.5 μg total RNA using oilgodT primers and RevertAid Reverse Transcriptase (Thermofisher). Quantitative reverse transcriptase PCR (qRT-PCR) was performed in a 384-well plate using cDNA (3 ng), gene specific primers (250 nM, list provided in Supplemental file), and iTaq Universal SYBR Green Supermix (1X, Bio Rad). Relative gene expression was calculated using the delta CT or delta-delta CT method (Livak and Schmittgen, 2001). Template normalization was done using two housekeeping genes, 18S ribosomal RNA and Ubiquitin-conjugating enzyme *E2*. Gene accession numbers and primer sequence information are presented in Supplemental file 1.

### Micro Chromatin Immunoprecipitation (μChIP) assay from protoplasts

Transcription factor PdWND1B was cloned in-frame with 10X Myc tag and used to transfect protoplasts derived from *Populus* 717-1B4 tissue culture grown plants (Guo et al, 2012). ChIP assays were performed using the modified protocol from Dahl and Collas (2008) and Adli and Bernstein (2011). Briefly, transfected protoplasts were resuspended in W5 solution (154mM NaCl, 125 mM CaCl_2_, 5 mM KCl, 2 mM MES (pH 5.7)), crosslinked by adding 1% (v/v) formaldehyde and gently rotating the tubes for 8 min. To stop crosslinking, Glycine was added to a final concentration of 0.125 M and gently rotated at room temperature for 5 min. The crosslinked protoplasts were washed once with W5 solution and lysed by mixing with SDS Lysis Buffer (50 mM Tris-HCl (pH 8.0), 100 mM NaCl, 10 mM EDTA (pH 8.0), 1% SDS, 1 mM PMSF, protease inhibitor) followed by incubation on ice for 10 min with intermittent and brief vortexing. The lysate was supplemented with RIPA ChIP Buffer (10mM Tris-HCl (pH 7.5), 140 mM NaCl, 1 mM EDTA (pH 8.0), 1% Triton X-100, 0.1% Na-deoxycholate, 1 mM PMSF, protease inhibitor) and sonicated for 150 s with 0.7 s ‘On’ and 1.3 s ‘Off’ pulses at 20% power amplitude on ice using Branson 450 Digital sonifier to generate 150- to 600-bp chromatin fragments. Additional ice-cold RIPA ChIP buffer was added to aliquot the sample into three separate tubes – 500 μl Antibody (Ab) sample, 500 μl No-Antibody (NAb) sample and 75μl input chromatin. To the Ab sample, 0.75-1 μg anti-c-Myc antibody (Sigma-Aldrich #C3956) was added and gently rotated overnight at 4°C. Protein A Mag Sepharose (Sigma-Aldrich #28-9440-06) beads were washed with RIPA buffer (10 mM Tris-HCl (pH 7.5), 140 mM NaCl, 1 mM EDTA (pH 8.0), 1% Triton X-100, 0.1% SDS, 0.1% Na-deoxycholate), added to Ab and NAb samples and gently rotated at 4°C for 120 min. The beads were then collected, washed twice with low-salt wash buffer (150 mM NaCl, 0.1% SDS, 20 mM Tris-HCl (pH 8.0), 2 mM EDTA (pH 8.0), 1% Triton X-100), twice with LiCl buffer (0.25 M LiCl, 1% Na-deoxycholate, 10 mM Tris-HCl (pH 8.0), 1% NP-40, 1 mM EDTA (pH 8.0)) and twice with TE Buffer (10 mM Tris-HCl (pH 8.0), 1 mM EDTA (pH 8.0)). The beads were subjected to reverse crosslinking by adding Complete Elution Buffer (20 mM Tris-HCl (pH 8.0), 5 mM EDTA (pH 8.0), 50 mM NaCl, 1% SDS, 50 μg/ml Proteinase K) and incubating for 120 min on thermomixer at 68°C and 1300 rpm to elute protein-DNA complexes. Input samples were added with elution buffer (20 mM Tris-HCl (pH 8.0), 5 mM EDTA (pH 8.0), 50 mM NaCl) and 50 μg/ml Proteinase K before placing on thermomixer. After incubation, supernatants were collected and the ChIP DNA was purified using MinElute PCR Purification Kit (Qiagen #28004). Real-time PCR was performed for the ChIPed DNA by promoter specific primers (Supplemental File 1) and the obtained Ct values were used to calculate the signal intensity by Percent Input Method. At least three biological replicates (with two technical replicates each) representing independent protoplast transfections were used. The ChIPed DNA was also used for PCR reactions by promoter specific primers to analyze the products on agarose gel.

### Transcriptional activator assay

The coding sequence (CDS) of *WND1B* was in-frame cloned in a Gal4 binding domain (GD) effector vector (Wang et al, 2007). For the trans-activator assays, the GD-fusion constructs were co-transfected with Gal4:GUS reporter construct into *Populus* 717-1B4 protoplasts (Guo et al, 2012). For the trans-repressor assays, GD-fusion constructs were co-transfected with LexA binding-domain fused VP16 (LD-VP) and LexA:Gal4:GUS reporter (Wang et al, 2007). An empty GD effector vector was co-transfected with reporter vectors for the control experiments. The transfected protoplasts were incubated in dark for 16-20 h and GUS activity was quantitatively measured. All the protoplast transfections were included with equal amounts of 35S:Luciferase reporter and Luciferase activity was used for normalization of GUS activity.

### Cellulose and lignin estimation

Cellulose was estimated on debarked, ground, and air-dried stem tissue using the anthrone method (Updegraff, 1969). Stem sample (25 mg) was first digested with 500 μl of acetic - nitric acid reagent (100 ml of 80% acetic acid mixed with 10 ml of nitric acid) at 98°C for 30 min. After cooling, the sample was centrifuged, the supernatant was discarded, and the remainder was washed with water. After brief centrifugation, water was discarded, and the pellet was digested with 67% (v/v) sulfuric acid for 1 h at room temperature. An aliquot of the mix was diluted (1:10) with water. In a PCR tube, 10 μl of diluted reaction mix, 40 μl of water, and 100 μl of freshly prepared anthrone reagent (0.5 mg anthrone ml^−1^ of cold concentrated sulfuric acid) was added and heated for 10 min at 96°C. Samples were cooled and absorbance (A630) was measured. Cellulose was then estimated based on the absorbance of glucose standards. Lignin and its monomer composition was analyzed using pyrolysis molecular beam mass spectrometry at the National Renewable Energy Laboratory as described previously (Mielenz *et al*., 2009).

### Stem carbohydrate composition analysis

Roughly 25 mg of air-dried stem sample was weighed in a 2-ml tube and extracted twice at 85°C with a total of 2 ml of 80% ethanol. The supernatant was collected in a new 2-ml tube and was re-extracted with 50 mg activated charcoal (Sigma) to eliminate pigments that interfere with sugar analysis. A 1 ml aliquot of the pigment free extract was incubated overnight in a heating block maintained at 50°C and the resulting pellet was dissolved in 120 μl of water. A 10 μl aliquot was used for estimation of sucrose and glucose using assay kits (Sigma). Starch from the pellet was digested using 1U of *α-amylase* (from *Aspergillus oryzae*, Sigma) and amyloglucosidase (from *Aspergillus niger*, Sigma). After starch removal, the pellet was dried overnight at 95°C and used for estimating structural sugars. Roughly, 5 mg of sample was weighed in a 2-ml tube and digested with 50 μl of 75% v/v H_2_SO_4_ for 60 min. The reaction was diluted by adding 1.4ml of water, tubes were sealed using lid-locks, and autoclaved for 60 min in a liquid cycle. After cooling, the sample was neutralized with CaCO_3_ and sugar composition was estimated using high performance liquid chromatography (HPLC, LaChrom Elite^®^ system, Hitachi High Technologies America, Inc.) as described previously (Fu *et al*., 2011; Yee *et al*., 2012).

### Glucose release and ethanol conversion

Separate hydrolysis and fermentation (SHF) was used to evaluate digestibility of biomass samples as described previously (Fu *et al*., 2011; Yee *et al*., 2012). Extract free biomass was autoclaved for sterilization purposes and the hydrolysis and fermentations were performed in biological triplicate at 5.0% (w/v) biomass loading in a total volume of 20 ml at a pH of 4.8 with a final concentration of 50 mM citrate buffer and 0.063 mg ml^−1^ streptomycin. The hydrolysis was performed using commercial hydrolytic enzyme blends (Novozymes, Wilmington, DE, USA). Cellic^®^-Ctec2 was loaded at 20 mg protein gram^−1^ dry biomass, and Novozyme 188 and Cellic^®^ Htec2 were loaded at 25% and 20% (v/v) of Ctec2, respectively. The biomass and enzymes were incubated at 50°C and 120rpm for 5 days. The hydrolysate was then fermented with *Saccharomyces cerevisiae* D5A (ATCC 200062) at 35°C and 150 rpm with a final concentration of 0.5% (w/v) yeast extract. Hydrolysate and fermentation broth samples were analyzed for glucose and ethanol using HPLC equipped with a refractive index detector (model L-2490). The products were separated on an Aminex^®^ HPX-87H column (Bio-Rad Laboratories, Inc.) at a flow rate of 0.5 ml min^−1^ of 5.0 mM sulfuric acid and a column temperature of 60°C and were quantified as described previously (Fu *et al*., 2011; Yee *et al*., 2012).

## Results and Discussion

### Phylogenetic analysis, gene expression, and localization

In *Arabidopsis*, at least three NAC transcription factors, *SND1, NST1*, and *VND7* have a proposed role in regulation of secondary cell wall formation. To retrieve their sequence orthologs from *Populus trichocarpa*, protein sequences of the three genes were blasted in *Phytozome (v 9.1*) and the two best hits were retrieved for each sequence resulting in a total of six sequences. These were designated as *PtrWND1A* (Potri.011G153300), *PtrWND1B* (Potri.001G448400), *PtrWND2A* (Potri.014G104800), *WND2B* (Potri.002G178700), *PtrWND6A* (Potri.013G113100) and *PtrWND6B* (Potri.019G083600). The nomenclature used in this study was based on Zhong et al., (2010). All six genes have alternate names; *WND1B* has also been named *SND1A2* or *VNS11* in previously reports (Li *et al*., 2012; Ohtani *et al*., 2011). In the phylogenetic tree developed using protein sequences, *PtrWND1A* and *PtrWND1B* were clustered together and share 94% similarity at the protein level (Figure 1; Supplemental file 2), suggesting they originated from a recent genome duplication (Tuskan *et al*., 2006). They share only approximately 50% similarity with *AtSND1* and *AtNST1*, approximately 56% with *PtrWND2A* and *PtrWND2B*, and approximately 41% with *PtrWND6A* and *PtrWND6B*, but more than 83% with *RcNAC* (*Ricinus communis*) and *JcNAC013* (*Jatropha curcas*). *PtrWND2A* and *PtrWND2B* are clustered together and share 88% similarity, while *PtrWND6A* and *PtrWND6B* share 92% similarity. Protein sequence alignment revealed they are highly conserved in the NAC domain located in the N terminal region. Conversely, they are highly diverse in the C terminal region, which has putative activation domains (Olsen *et al*., 2005; Xie *et al*., 2000). At least 163 NAC domain transcription factors have been reported in *Populus*. Based on phylogenetic analysis, these are classified into 18 groups (Hu *et al*., 2010). *PtrWND1A* and *B* and *PtrWND2A* and *B* are closely clustered in the NAC-B subgroup, while *PtrWND6A* and *B* are clustered in the NAC-O subgroup.

**Figure 1.**
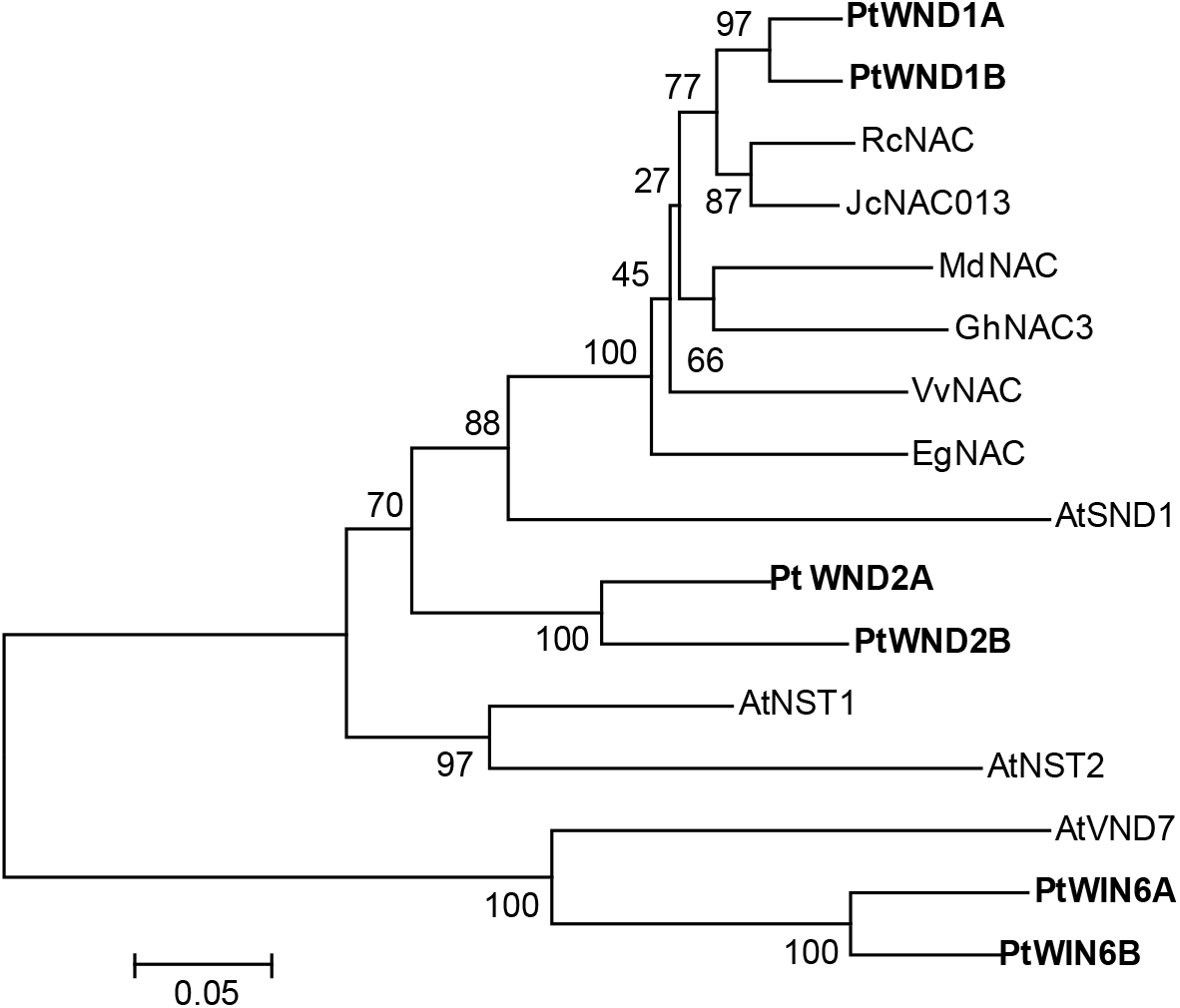
Phylogenetic analysis of selected secondary cell wall associated NAC transcription factors from *Populus* and other plant species. Transcription factors from *Populus* are in bold. The percentage of replicate trees in which the associated taxa clustered together in the bootstrap test (500 replicates) are shown next to the branches. Accessions are provided below. AtSND1: At1g32770 (*Arabidopsis thaliana*); AtNST1: At2g46770; AtNST2: At3g61910; AtVND7: AT1G71930; RcNAC: XP_002518924 (*Ricinus communis*); VvNAC: XP_002279545 (*Vitis vinifera*); JcNAC013: AGL39669 (*Jatropha curcas*); MdNAC: NP_001280877 (*Malus domestica*), GhNAC3: ADN39415 (*Gossypium hirsutum*); EgNAC: KCW72583 (*Eucalyptus grandis*). PtWND1A (Potri.011G153300), PtWND1B (Potri.001G448400), PtWND2A (Ptri.014G104800), WND2B (Potri.002G178700), PtWND6A (Potri.013G113100) and PtWND6B (Potri.019G083600).

The expression of the above six NAC transcription factors was studied in eight different tissues of *Populus deltoides* including YL (young leaf), ML (mature leaf), YS (young stem), MS (mature stem), PH (phloem), XY (xylem), RT (root), and PT (petiole). In *Populus*, *WND1B* undergoes alternate splicing, resulting in two variants designated as the small and large variants. The large variant retains intron 2 (Li *et al*., 2012; Zhao *et al*., 2014). In this study, to account for both splice variants, primers were designed in the region common to both variants. In general expression of all the genes was much higher in xylem than in other tissues (Supplemental file 3). Within xylem, expression of *PdWND1A* and *PdWND1B* was much higher relative to the other genes. Among other tissues, expression of *PdWND1A* was much higher relative to the other genes except in phloem, where *PdWIN2B* was strongly expressed. Expression of *PdWIN6A* and *PdWIN6B* was weaker in all tissues relative to the other genes. In *Populus*, these transcription factors are most abundantly expressed in stems. *In situ* localization studies suggested both *PtrWND1B* and *PtWND6A* are expressed in xylem vessels and fibers and in phloem fibers after secondary growth, whereas in primary xylem vessels only *PtrWND6A* expression was observed, suggesting developmental regulation (Ohtani *et al*., 2011). The two abundantly expressed genes, *PdWND1A* and *PdWND1B*, were selected for localization studies using tobacco infiltration. *GFP:PdWND1A* and GFP: *WND1B* were colocalized with DAPI stain confirming both *PdWND1A* and *PdWND1B* are targeted to the nucleus (Supplemental file 4). *AtSND1* and *PtWND1B* have previously been localized to the nucleus supporting their function as transcription factors (Li *et al*., 2012; Zhong *et al*., 2006).

### Plant morphology and growth

In the present study, we focused on studying the functional aspects of *PdWND1B* through over-expression and RNAi-mediated suppression using a xylem specific *4CL* promoter. In order to selectively down-regulate *PdWND1B*, sequence in the 3′ region that has distinct differences with *PdWND1A* was selected for RNAi construct development. In our preliminary study, six independent over-expression (OE) and six independent RNAi lines were propagated in the greenhouse. Plant height of over-expression lines was not different as compared to controls, but RNAi lines were shorter (Supplemental file 5A). Lignin content was significantly higher in all over-expression lines but showed a slight decreasing trend in RNAi lines (Supplemental file 5B). In-depth characterization was performed on two to four selected lines and data presented in this study is representative of two over-expression lines (designated as OE2 to OE4) and two RNAi suppression lines (Ri1 and Ri4). The extent of alteration in *PdWND1B* expression in transgenic lines was measured using qRT-PCR. As compared to control lines, *PdWND1B* expression was increased by 40-fold in OE4 and by 23-fold in OE2 (Figure 2). In RNAi lines, *PdWND1B* expression was reduced by 73% in Ri4 and by 65% in Ri1.

**Figure 2.**
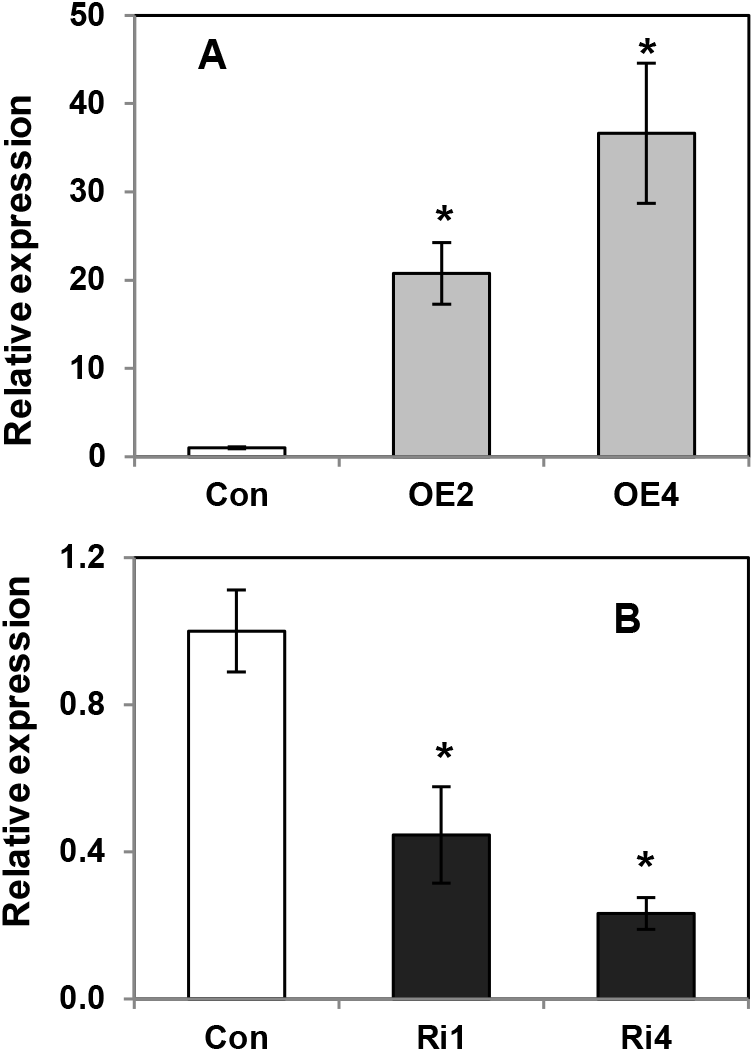
Relative expression of *PdWND1B* in control and transgenic lines. Gene expression (arbitrary units) in control (Con), over-expression (OE) lines (A), and RNAi suppressed (Ri) lines (B) was relative to the housekeeping genes *Ubiquitin conjugating enzyme E2* and *18S RNA*. The data represents means ± SE (n = 3). * Indicates statistical significance based on Student’s *t*-test (p ≤ 0.05).

At the time of harvest (~six months of growth), control plants reached an average of 130 cm (Figure 3A). The OE plants were similar in height with that of controls. However, Ri lines were significantly shorter by 40 to 50% and reached an average of 66 to 78 cm (Figure 3A). A similar trend was also observed in stem diameter. As compared to controls, stem diameter in OE expression lines was not significantly altered but was reduced by 40% in Ri1 and Ri4 (Figure 3B). The combined effect of reduced plant height and stem diameter resulted in a roughly 75% reduction in total stem dry weight in Ri1 and Ri4 lines (Figure 3C). RNAi lines also developed smaller leaves and thus had a roughly 70% reduction in leaf weight (Figure 3D).

**Figure 3.**
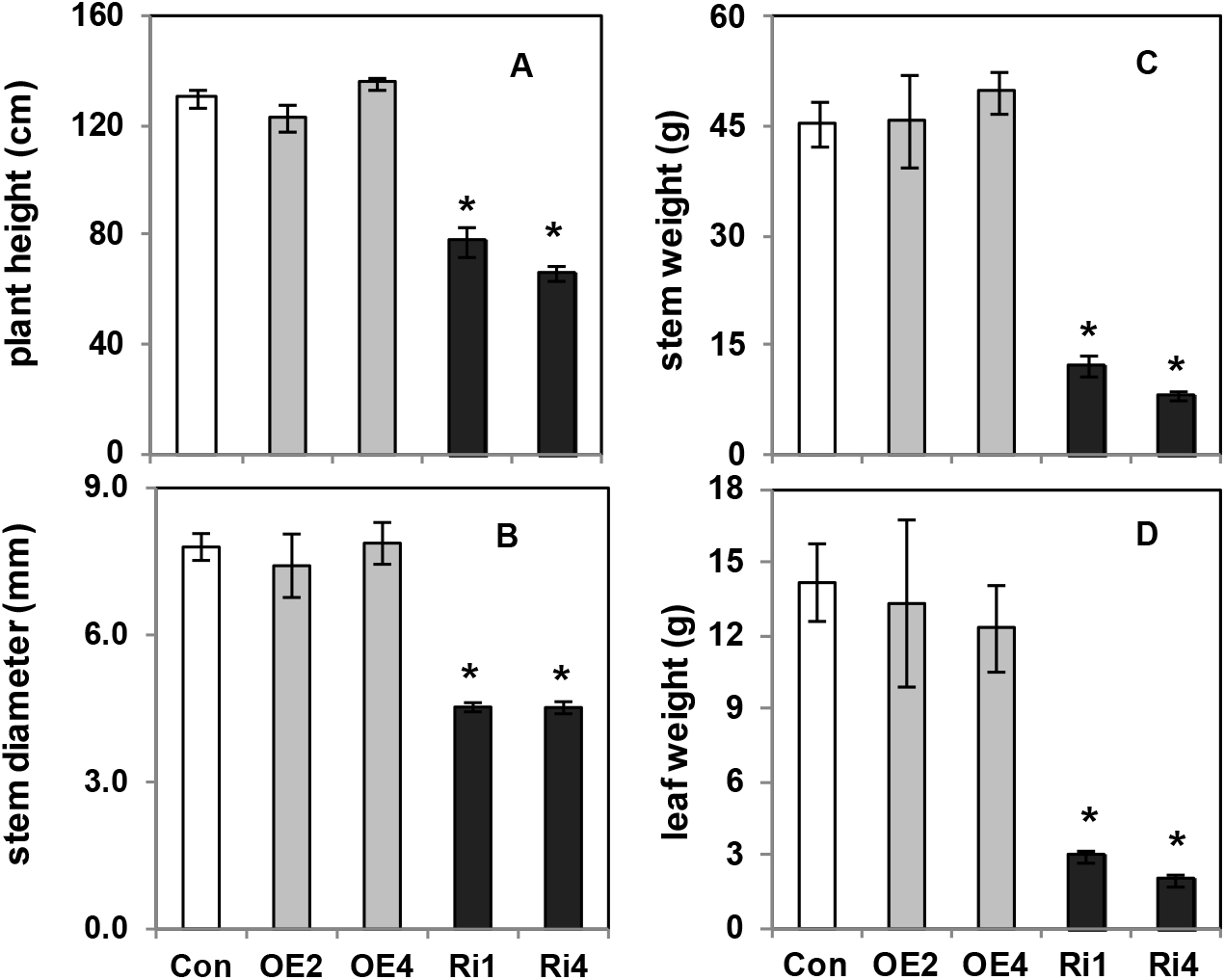
Growth and biomass productivity in control and *PdWND1B* transgenic lines. Plant height (A), stem diameter (B), stem weight (C), leaf weight (D) of empty vector transformed control (Con), and PdWND1B over-expression (OE) and RNAi suppressed (Ri) lines. The data represents means ± SE (n = 3). * Indicates statistical significance based on Student’s *t*-test (p ≤ 0.05).

Evidence suggests that *SND/WND* are required for normal plant development (Zhao *et al*., 2014; Zhong *et al*., 2010). Over-expression of the full-length coding region of *AtSND1* in *Arabidopsis* and *PtWIN2B* or *PtWIN6B* in *Populus tremula x alba*, under the control of a CaMV 35S promoter, resulted in plants with weaker stems, small leaves, and stunted growth. This strongly supports the hypothesis that the WNDs play a significant role in maintenance of growth and development (Zhong *et al*., 2006; Zhong *et al*., 2011). In contrast, over-expression of the *PtWND1B* whole gene (including exons and introns) in *Populus x euramericana*, under the control of a CaMV 35S promoter, did not affect plant growth, but only reduced leaf size (Zhao *et al*., 2014). Our study included overexpression of the shorter variant of *PdWND1B* under the control of a xylem-specific promoter and the observation of no apparent growth impact in overexpression lines. Previous study reported that over-expression of the *PtWND1B* longer splice variant in *Populus x euramericana*, under the control of its own promoter, affected plant development, but the same effect was not observed when the small variant of *PtWND1B* was over-expressed (Zhao *et al*., 2014). Zhong *et al*., (2006) also report that an *Atsnd1* mutation did not affect plant development However, consistent with our study in *Populus*, down-regulation of *PtWND1B*, controlled by its own promoter, resulted in plants with weak stems that did not grow straight (Zhao *et al*., 2014). Therefore, it appeared that *WND* genes may have species-specific effect on plant growth and development. It is also possible that the differences in promoters used (i.e., native promoter (Zhao *et al*., 2014) and tissue-specific promoter (in the present study) may contribute to differences in phenotypic observations between *Arabidopsis* and *Populus*.

### Structural polymers

WND transcription factors have a proposed function in secondary cell wall biosynthesis. Therefore, the effect of altered *PdWND1B* expression on secondary cell wall composition were studied in stems. Stem secondary cell walls are composed predominantly of cellulose, lignin, and hemicellulose (Bailey, 1938; Darvill *et al*., 1980). Cellulose, estimated by the anthrone method, was significantly reduced by 9 to 13% in OE lines, but was increased by 6% in RNAi lines (Figure 4A). Lignin content was significantly increased in OE lines but decreased in RNAi4 (Figure 4B). To understand changes in other sugars, stem cell walls were digested and sugars were quantified using HPLC. Glucose and xylose were the predominant sugars in control plant stem material, at 45% and 15%, respectively (Figure 5). However, while glucose was reduced in OE lines, xylose, representing the hemicellulose fraction, was significantly increased (Figure 5). Levels of minor sugars including galactose, arabinose, and mannose were also significantly reduced in OE lines. Trace compounds, 5-(Hydroxymethyl) furfural was reduced (up to four fold) in RNAi lines, while 2-furfural was significantly reduced by 60 to 75% in RNAi lines.

**Figure 4.**
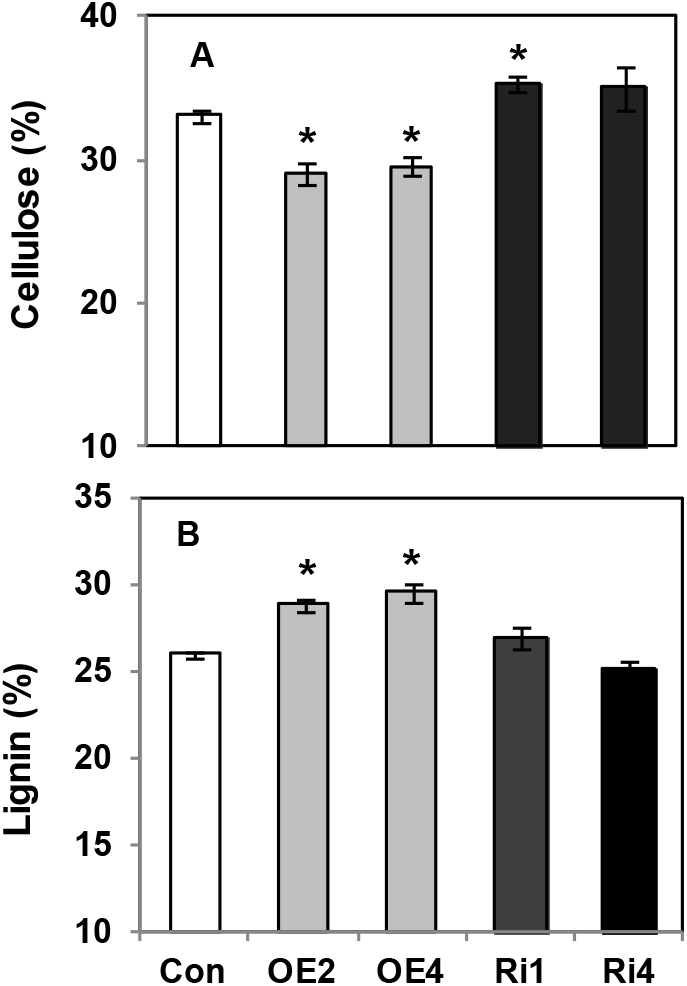
Stem cell wall composition of control and *PdWND1B* transgenic lines. Levels of cellulose (A) and lignin (B) in empty vector transformed control (Con), *PdWND1B* over-expression (OE), and RNAi suppressed (Ri) lines. The data represents means ± SE (n = 3 to 5). * Indicates statistical significance based on Student’s *t*-test (p ≤ 0.05).

**Figure 5.**
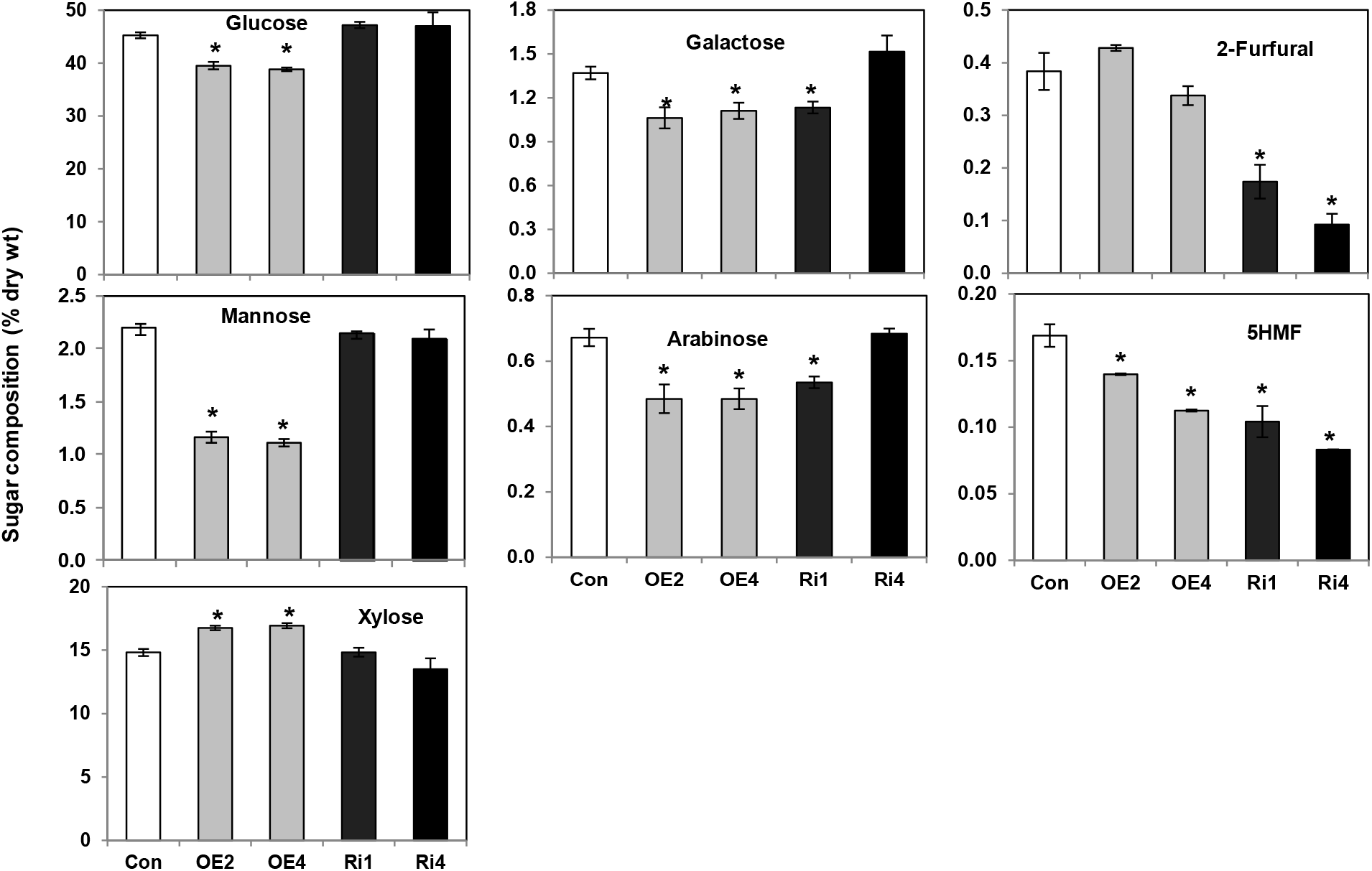
Sugar composition in stem cell walls of control and *PdWND1B* transgenic lines. Levels of different sugars in empty vector transformed control (Con), *PdWND1B* over-expression (OE), and RNAi suppressed (Ri) lines. The data represents means ± SE (n = 3). * Indicates statistical significance based on Student’s *t*-test (p ≤ 0.05).

Over-expression of *AtSND1* induced ectopic deposition of lignified secondary cell walls in leaf and stem epidermal and mesophyll cells that normally do not undergo lignification (Zhong *et al*., 2006). In addition, cellulose and hemicellulose were also deposited. A similar response was observed in *Populus*, where *PtWND2B* and *PtWND6B* were over-expressed under the control of a CaMV 35S promoter (Zhong *et al*., 2011). To address the biomass chemistry context of the present study, over-expression of *PdWND1B* in our study was driven by a xylem-specific promoter to avoid confounding growth effects arising from ectopic lignification. In the context of stem cell wall phenotype, our results indicate an increase in lignin and xylose in stems of OE lines while cellulose levels were reduced. A negative relationship has been proposed between levels of cellulose and lignin (Hu *et al*., 1999). We observed an increase in lignin and a concomitant decrease in cellulose of overexpression lines relative to the control. In *Arabidopsis*, silencing of *AtSND1* and *AtNST1* simultaneously reduced lignin, cellulose, and hemicellulose (Zhong *et al*., 2007a). In the present study, significant differences were not observed in levels of lignin or other sugars in RNAi lines, suggesting that the reduction in expression and function of *PdWND1B* potentially is partly compensated by other members of the NAC family (i.e., PdWND1A) members. In future studies, it would be interesting to generate and characterize double knockout/knockdown plants of *PdWND1A* and *PdWND1B*, and similarly, for other closely related paralogs, which can address the potential functional redundancy and reveal their more precise functions in secondary cell wall biosynthesis.

### Sugar release and ethanol conversion

The effect of altered cell wall composition on sugar release and ethanol conversion was studied in OE and RNAi lines. Glucose release was significantly reduced by 65 to 70% in OE plants compared to that of control plants (Figure 6A). This is consistent with a significant reduction in ethanol production from biomass. In contrast, glucose release was increased by 15% and 20% in RNAi1 and RNAi4 lines, respectively; however, ethanol production was increased (30%) only in RNAi4, the lines with greater downregulation (Figure 6B).

**Figure 6.**
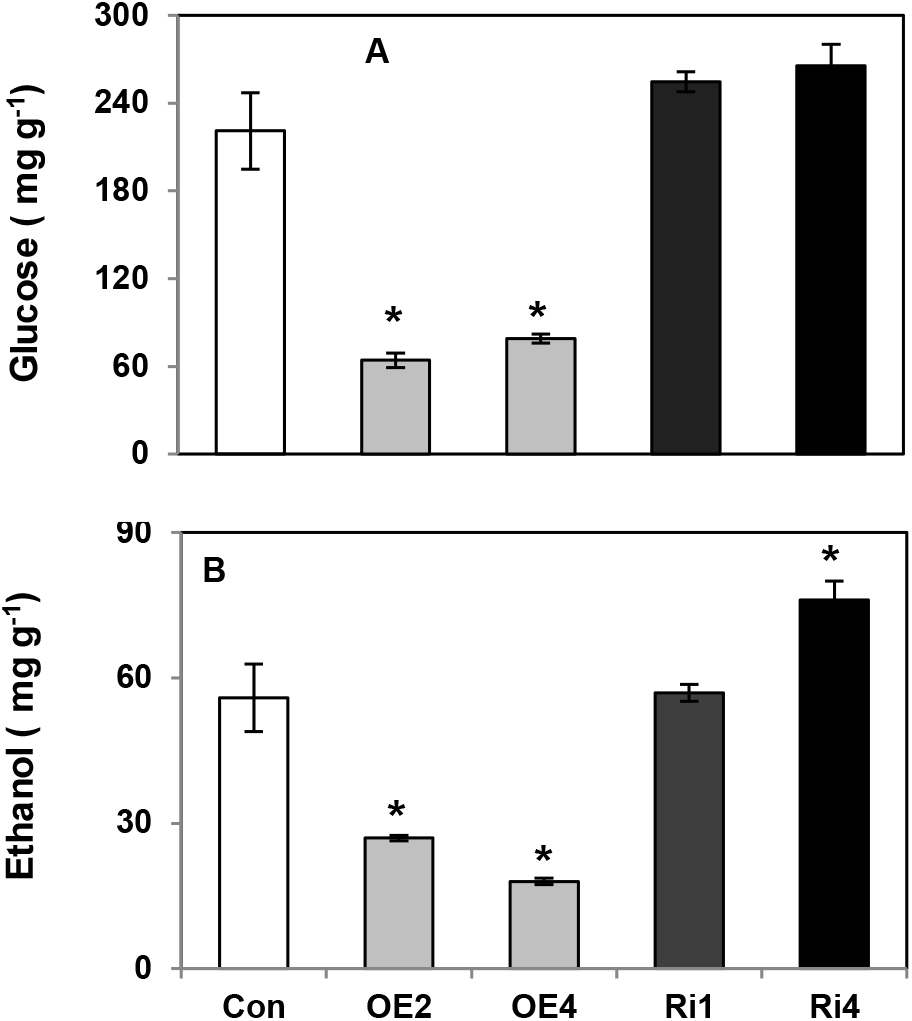
Glucose release and ethanol conversion efficiency from stems of control and *PdWND1B* transgenic lines. Levels of glucose (A) and ethanol (B) in empty vector transformed control (Con), *PdWND1B* over-expression (OE), and RNAi suppressed (Ri) lines. The data represents means ± SE (n = 3). * Indicates statistical significance based on Student’s *t*-test (p ≤ 0.05).

Biomass recalcitrance is determined by many parameters, but predominantly by cellulose and lignin content and composition. Lignin content and S:G ratio have been reported to influence sugar release efficiency in poplar (Studer *et al*., 2011). An increase in lignin content and decrease in cellulose content had a strong negative impact on sugar release efficiency and ethanol conversion in OE lines in this study.

### Gene expression changes

In *Arabidopsis* and *Populus*, over-expression of *AtSND1* and *PtrWND2B* induced expression of a cascade of other transcription factors and structural genes involved in lignin, cellulose, and hemicellulose formation (Zhong *et al*., 2006; Zhong *et al*., 2011). A set of 26 *Populus* transcription factors homologous to *Arabidopsis* secondary cell wall associated transcription factors induced by *AtSND1* over-expression were studied here. The expression of all 26 transcription factors was examined in xylem cDNA libraries obtained from two OE lines and two RNAi lines. Over-expression of *PdWND1B* significantly increased expression of several MYBs. Among these, the most prominent were *NAC154, NAC156, MYB18, MYB75, MYB199, MYB167, MYB175, MYB28, MYB31* and *MYB189*, where the expression was increased by 3 to 9-fold (Figure 7A). However, the expression of two genes, *WIN2A* and *MYB165* was decreased by up to 65% in the same OE lines. In *PdWND1B* RNAi lines, expression of *WIN2A, MYB18, MYB152*, and *MYB175* were increased by 2- to 3-fold while that of *WIN2B, MYB2*, and *MYB161* were reduced by 60 to 80% compared to controls (Figure 7B).

**Figure 7.**
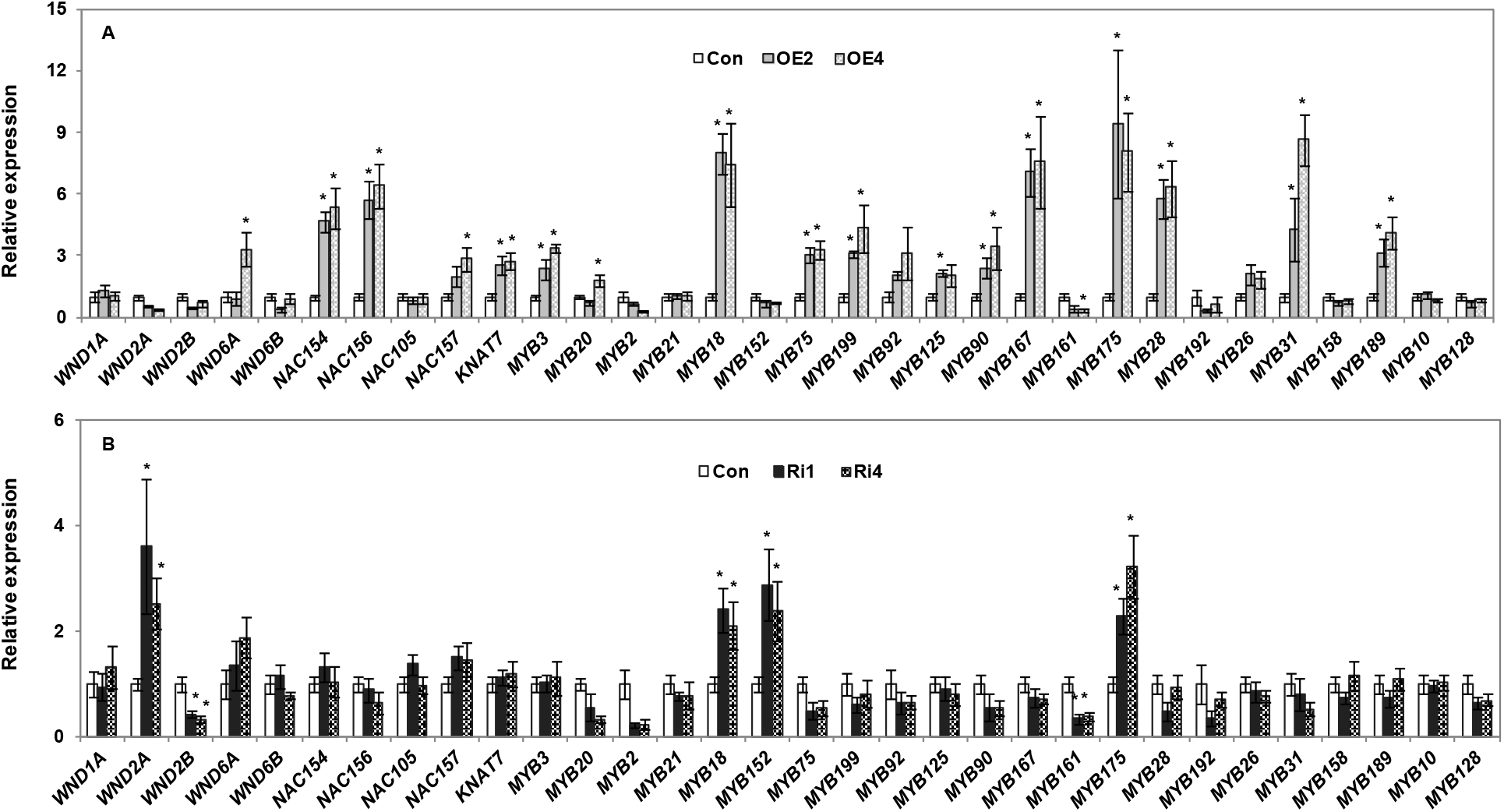
Expression of secondary cell wall related transcription factors in control and *PdWND1B* transgenic lines. Relative gene expression (arbitrary units) in control (Con), over-expression (OE) lines (A), and RNAi suppressed (Ri) lines (B) was calculated based on the expression of target genes relative to house-keeping genes, *Ubiquitin conjugating enzyme E2* and *18S RNA*, and then normalized to control. The data represents means ± SE (n = 3). * indicates statistically significant, *p* <0.05 based on Student’s *t*-tests.

In a previous study, over-expression of *PtrWND2B* induced expression of *PtrWND1A* and *B*, *PtrWND2A*, and *PtrWND6A* and B (Zhong *et al*., 2011). However, over-expression of *PdWND1B* induced only *PdWND6A* in our study. Also, *PtrWND2B* induced expression of all transcription factors except *PtrMYB152* (Wang et al., 2014). In contrast, several transcription factors were not induced in our study, suggesting that *WND1B* and *WND2B* may have distinct targets with some overlap. Alternatively, in the previous study, gene expression was quantified in leaf tissue where secondary wall formation is uncommon, while our study employed developing xylem tissue where secondary cell wall biosynthesis-related genes are viewed to be more specifically regulated by those TFs. The increase in *PdWND2A* in the *PdWND1B s*uppression lines indicates the existence of a compensatory mechanism. Although induction of *PdWND2A* or, more likely, other MYBs compensated for cell wall composition, they did not compensate and maintain normal growth in RNAi lines. In herbaceous plants such as *Arabidopsis*, *snd1* or *nst1* single mutants had no obvious growth defects, but *snd1 nst1* double mutants had severely affected stem strength suggesting that either one is sufficient for proper growth (Zhong *et al*., 2007a). In *Arabidopsis*, over-expression of *AtSND1* induced expression of *AtMYB46* (Zhong *et al*., 2007b), but over-expression of *PdWND1B* did not induce expression of *PdMYB002* and *PdMYB021*, the homologs of *AtMYB46*, implying the existence of potential species-specific regulation. Over-expression of *AtSND1* and *AtNST1* induced expression of *AtMYB58*. However, only *AtNST1* induced *AtMYB63* (Zhou *et al*., 2009). Our results were consistent with *Arabidopsis* in that over-expression of *PdWND1B* induced expression of *PdMYB28*, the closest homolog of *AtMYB58* but not *PdMYB192*, the closest homolog of *AtMYB63*, suggesting that WND/NAC master regulators have both redundant and distinct gene targets, and exhibit species-specificity in downstream regulation. *AtMYB58* and *AtMYB63* induced lignin formation but not cellulose and hemicellulose formation, suggesting that individual MYBs are specific to each pathway (Zhou *et al*., 2009). Relative to *PdWND1B* RNAi lines, the observed greater impact of *PdWND1B* overexpression on expression of cell wall transcription factor genes was also observed on expression of secondary cell wall (Shi et al. 2021) and sugar metabolism related genes (Figure 8).

**Figure 8.**
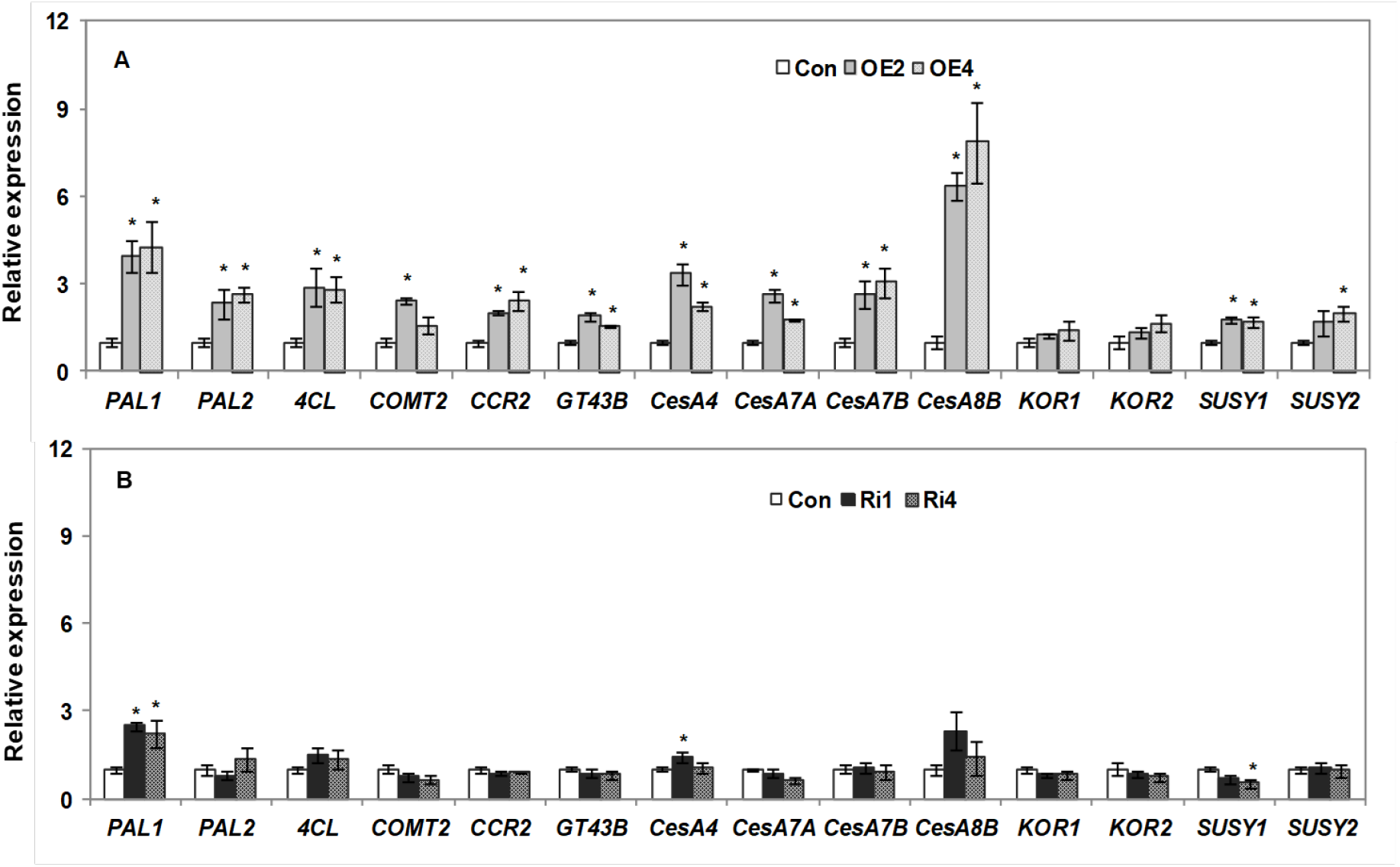
Expression of secondary cell wall related and sugar metabolism related genes in control and *PdWND1B* transgenic lines. Relative gene expression (arbitrary units) in control (Con), over-expression (OE) lines (A), and RNAi suppressed (Ri) lines (B) was calculated based on the expression of target genes relative to house-keeping genes, *Ubiquitin conjugating enzyme E2* and *18S RNA*, and then normalized to control. *PAL*, phenylalanine ammonia lyase; *4CL*, 4-coumarate:CoA ligase; *COMT*, caffeic acid/5-hydroxyconiferaldehyde O-methyltransferase; *CCR*, cinnamoyl-CoA reductase; *GT43*, Glucosyltransferase family 43; *CesA*, Cellulose synthase; *KOR*, Korrigan; *SUSY*, sucrose synthase. The data represents means ± SE (n = 3). * indicates statistically significant,*p* <0.05 based on Student’s *t*-tests.

### Promoter binding and transcriptional activation

PdWND1B has been previously reported as a transcription activator and is found to bind to promoters of MYB002 (Lin et al. 2013), as well as the newly reported cell wall transcriptional regulators, *HB3* (Badmi et al. 2018) and *EPSP* (Xie et al. 2018), in *Populus*. Transactivation assays confirmed that PdWND1B acts as a transcriptional activator and not as a transcriptional repressor (Supplemental File 5). *In vivo* DNA binding assay using micro-chromatin immunoprecipitation (μChIP) confirmed the binding of PdWND1B on the promoter of *PdMYB002*, a known target of Ptr-SND1-B1 (Lin et al., 2013) (Supplemental File 6), pointing to the overlapping functions of two poplar NAC homologs. Overexpression of PdWND1B induces the expression of a gene encoding 5-Enolpyruvylshikimate 3-Phosphate Synthase (EPSP), an enzyme that has been demonstrated activity as a transcriptional repressor and is involved in lignin biosynthesis (Xie et al., 2018). ChIP and transactivation assays suggest that PdWND1B binds to the promoters of the two *Populus EPSP* homologs, *EPSP1* and *EPSP2*, and activates their transcription *in vivo* (Supplemental File 7). These results indicate that PdWND1B is the upstream regulator of EPSP in lignin biosynthesis. The HD-ZIP III family of transcription factors have known roles in stem development (Robischon et al., 2011; Zhu et al., 2013). PdWND1B binds to the two homologs of the HD-ZIP III family of transcription factors, *PtHB3* and *PtHB4* and activates their transcription *in vivo* (Supplemental File 8). It has also previously been reported that PdWND1B binds to the promoter of a calmodulin binding protein *PdIQD10*, which is also involved in secondary cell wall biosynthesis (Badmi et al., 2018). Our results provide molecular evidence to further substantiate the role of PdWND1B as a master regulator of secondary cell wall biosynthesis during woody stem development in *P. deltoides*.

## Conclusion

Secondary cell wall composition depends on expression of *WND* transcription factors. The functional role of *WND1B* in *Populus* was studied by over-expression and down-regulation under the control of a xylem specific promoter. Over-expression of *PdWND1B* induced a cascade of transcription factors and structural genes involved in secondary cell wall biosynthesis. Phenotypic changes were aligned with molecular changes, specifically, over-expression of *PdWND1B* resulted increased lignin and xylose content, but decreased glucose resulting in a significant reduction in ethanol conversion. Down-regulation of *PdWND1B*, on the other hand, did not consistently alter lignin and cellulose content in stems but did impact other wall components and resulted in stunted growth. It is plausible that a functional compensation, as has been reported before, by other NAC members including *WND2A* and MYBs such as *MYB18, MYB152* and *MYB175*, in part explains the lack of significant impact on cell wall chemistry as a result of down-regulation of *PdWND1B*. Taken in total, our results suggest that *PdWND1B* does play a functional role in secondary cell wall biosynthesis through coordination with transcription factors and structural genes, which is further supported by the molecular evidence of its function to activate the transcription of several secondary cell wall pathway genes reported in the literature. In the future, studies designed to dissect the redundant and non-redundant functions of *PdWND1B*, its other homologs, and downstream transcription factors in stem, as well as root tissues, are needed to shed important and timely light on the redundant, conserved, and divergent mechanisms of plant biomass chemistry and productivity. Such fundamental understanding is critical to developing biodesign-based approaches to co-optimize aboveground performance for bio-derived fuels and products and soil health belowground.

## Abbreviations

4CL: *4*-coumarate-*CoA* ligase
DAPI: 4,6′-diamidino-2-phenylindole
CTAB: cetyltrimethylammonium bromide
HPLC: high performance liquid chromatography
LPI: leaf plastochron index
MEGA: Molecular Evolutionary Genetics Analysis
NAC: NAM, ATAF1/2 and CUC2
NST: NAC secondary wall thickening promoting factor
PBS: phosphate-buffered saline
qRT-PCR: quantitative reverse transcriptase
*PAL*: phenylalanine ammonia lyase
PCR: 
RNAi: RNA interference
S:G: syringyl to guaiacyl ratio
SHF: separate hydrolysis and fermentation
SHN: SHINE; SND, secondary wall associated NAC domains
VND: vascular related NAC domain
WND: wood associated NAC domain transcription factors
*4CL*: 4-coumarate:CoA ligase
*COMT*: caffeic acid/5-hydroxyconiferaldehyde O-methyltransferase
*CCR*: cinnamoyl-CoA reductase
*CesA*: Cellulose synthase
*KOR*: Korrigan
*GT43*: Glucosyltransferase family 43
*SUSY*: sucrose synthase

## Acknowledgements

We thank Brock Carter and Zackary Moore for inventory, propagation and maintenance of plants in ORNL greenhouses. This manuscript has been authored by UT-Battelle, LLC under Contract No. DE-AC05-00OR22725 with the U.S. Department of Energy. The United States Government retains and the publisher, by accepting the article for publication, acknowledges that the United States Government retains a non-exclusive, paid-up, irrevocable, world-wide license to publish or reproduce the published form of this manuscript, or allow others to do so, for United States Government purposes. The Department of Energy will provide public access to these results of federally sponsored research in accordance with the DOE Public Access Plan(http://energy.gov/downloads/doe-public-access-plan).

---

**Supplemental File 1.**
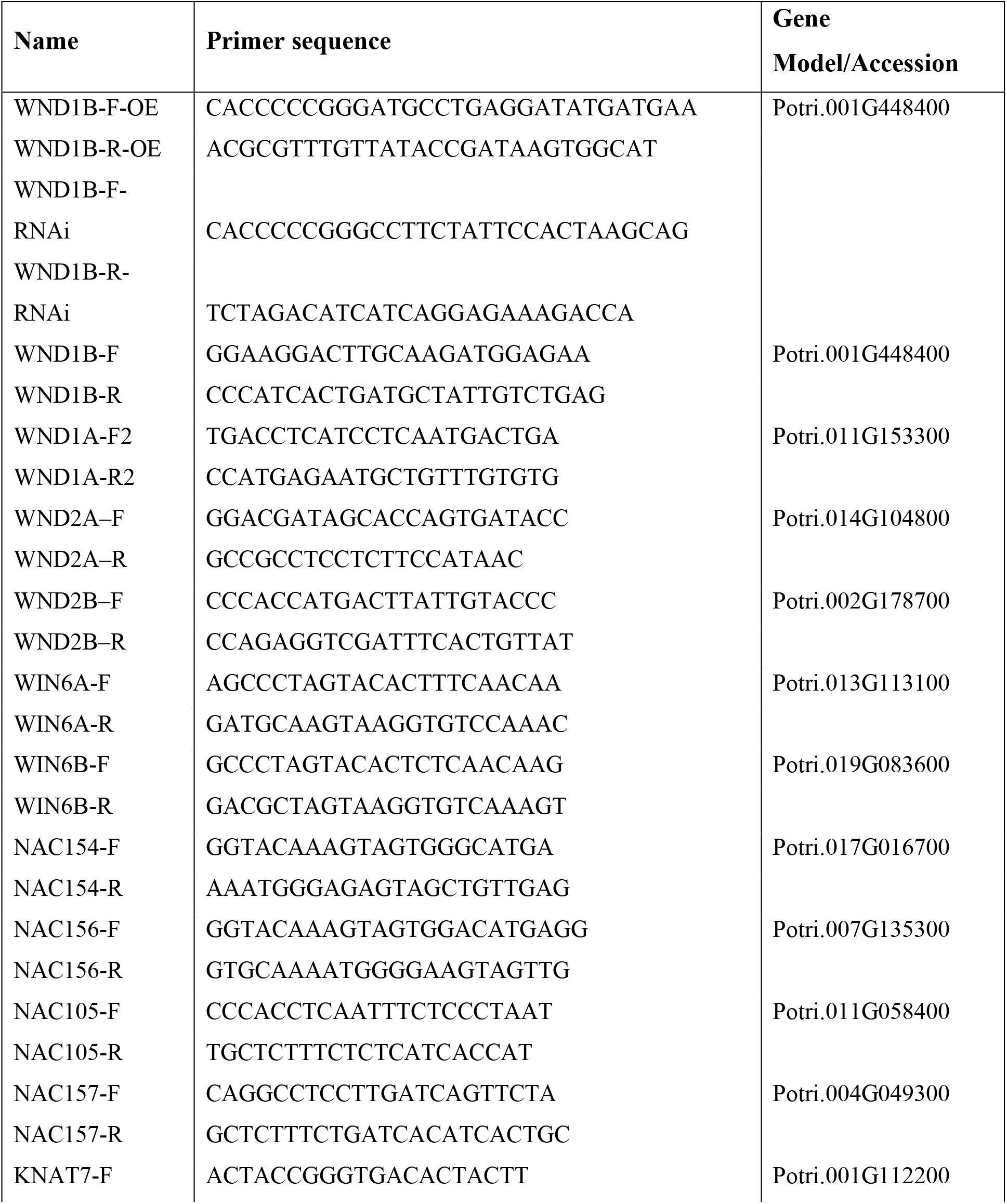
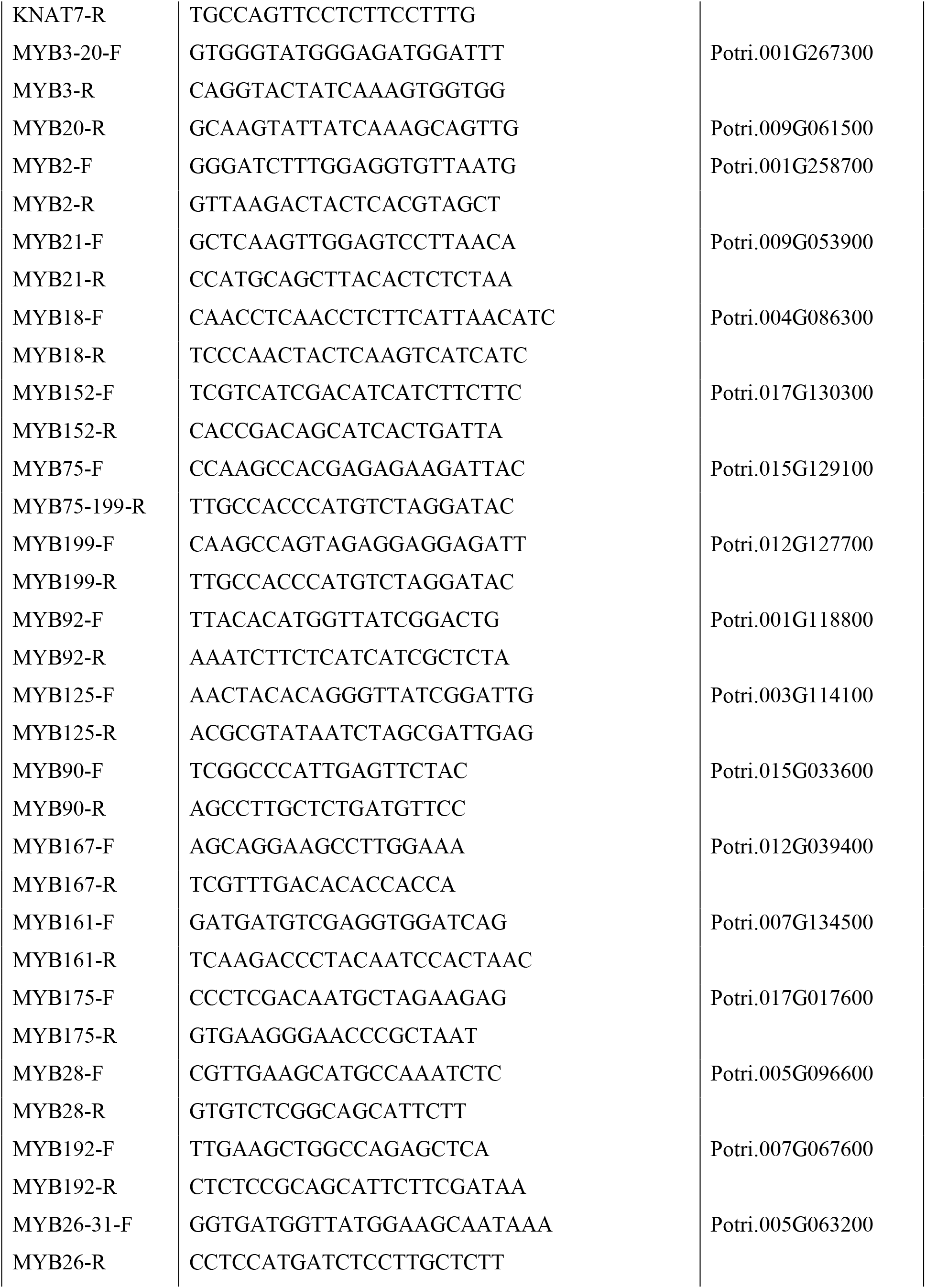
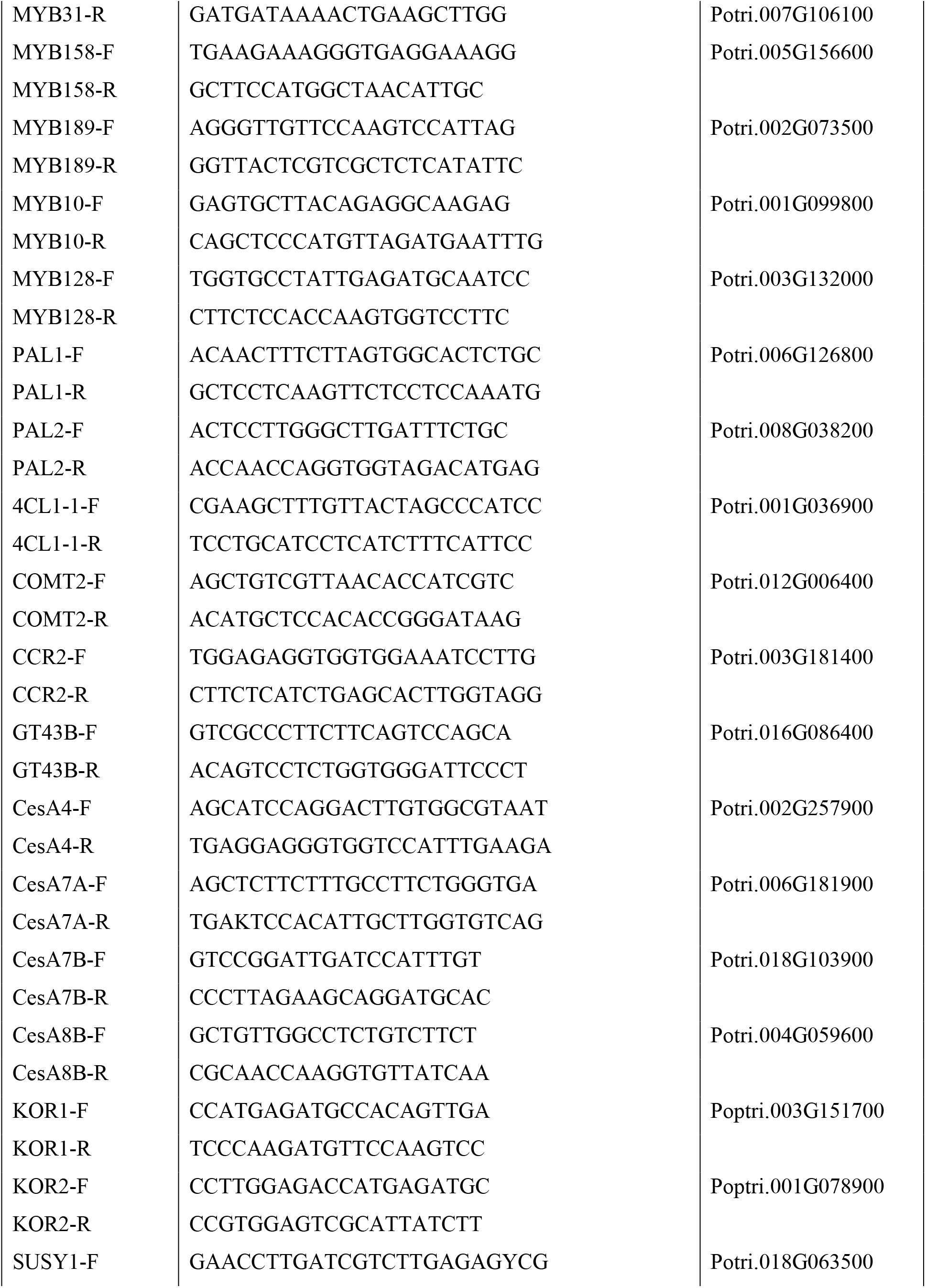
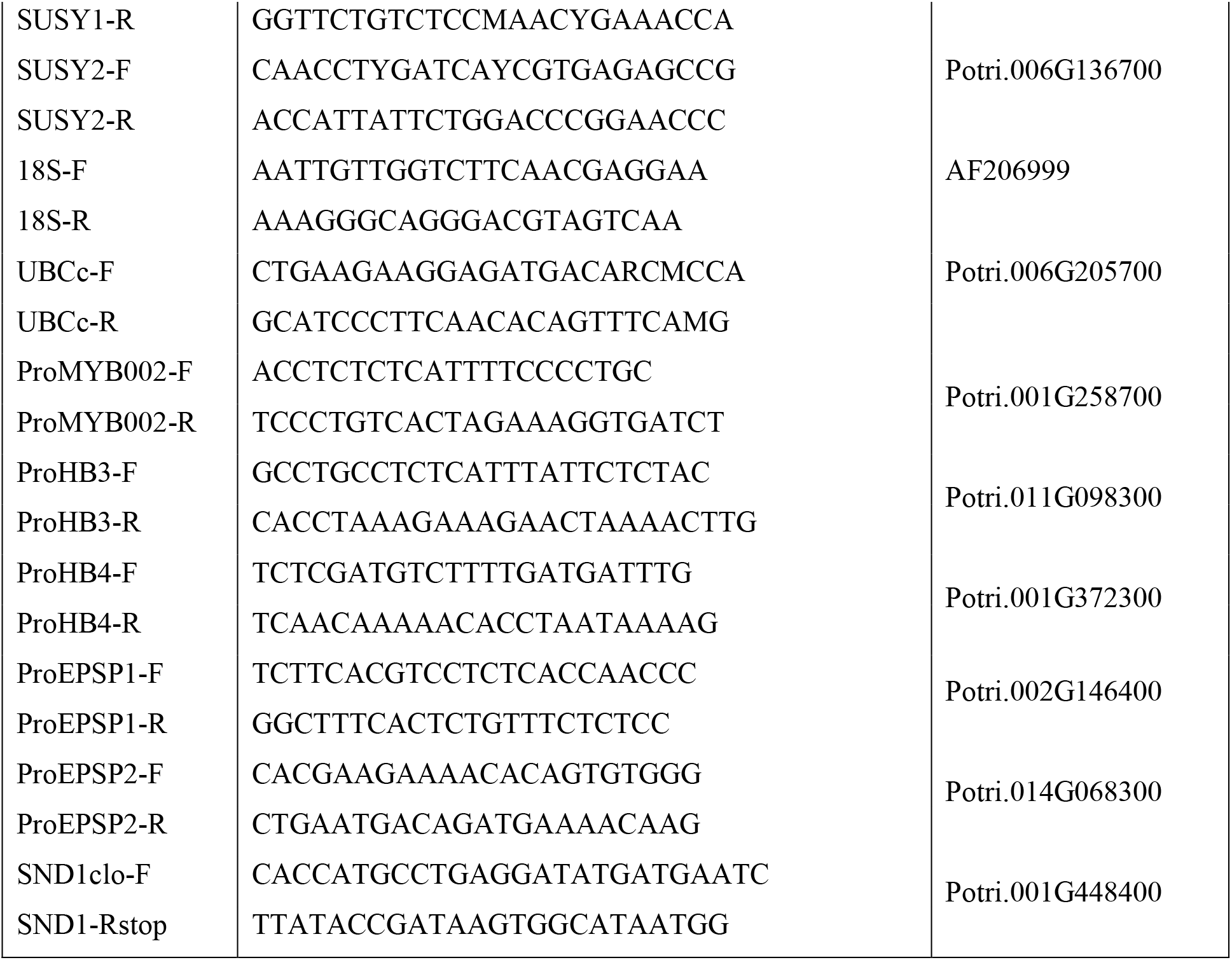
List of gene models and their primer sequence information. OE and RNAi primers were used to design over-expression and RNAi-knockout construct respectively and the rest for qRT-PCR. F and R indicate forward and reverse primers, respectively.

**Supplemental File 2.**
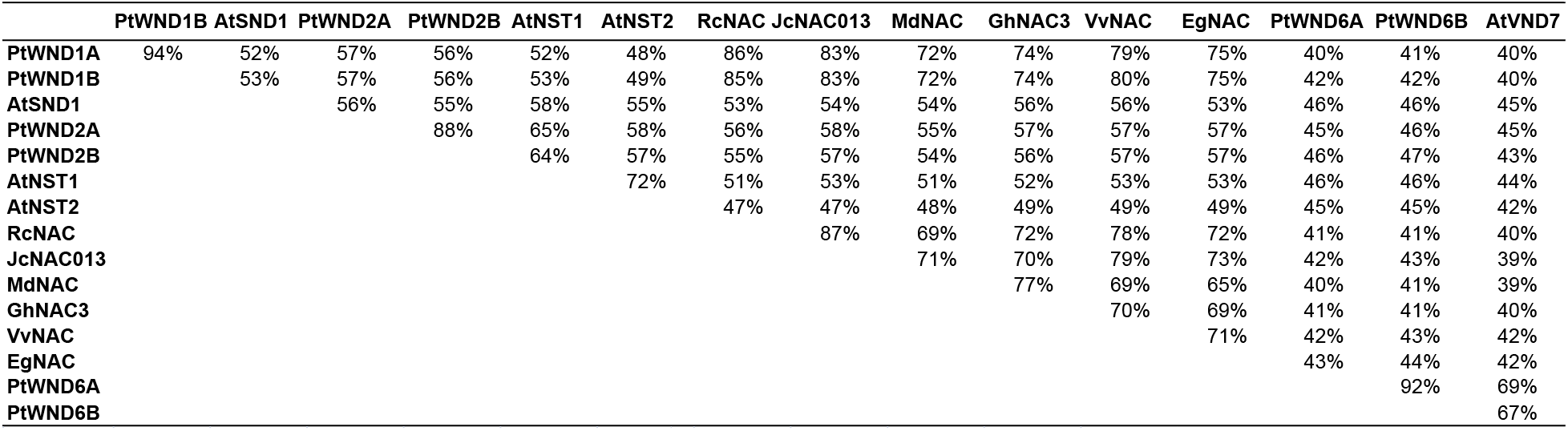
Percentage protein similarity matrix of selected secondary wall associated transcription factors from *Populus* and other species. Accessions are provided below. AtSND1: At1g32770 (*Arabidopsis thaliana*); AtNST1: At2g46770; AtNST2: At3g61910; AtVND7: AT1G71930; RcNAC: XP_002518924 (*Ricinus communis*); VvNAC: XP_002279545 (*Vitis vinifera*); JcNAC013: AGL39669 (*Jatropha curcas*); MdNAC: NP_001280877 (*Malus domestica*), GhNAC3: ADN39415 (*Gossypium hirsutum*); EgNAC: KCW72583 (*Eucalyptus grandis*). PtWND1A (Potri.011G153300), PtWND1B (Potri.001G448400), PtWND2A (Ptri.014G104800), WND2B (Potri.002G178700), PtWND6A (Potri.013G113100) and PtWND6B (Potri.019G083600).

**Supplemental File 3.**
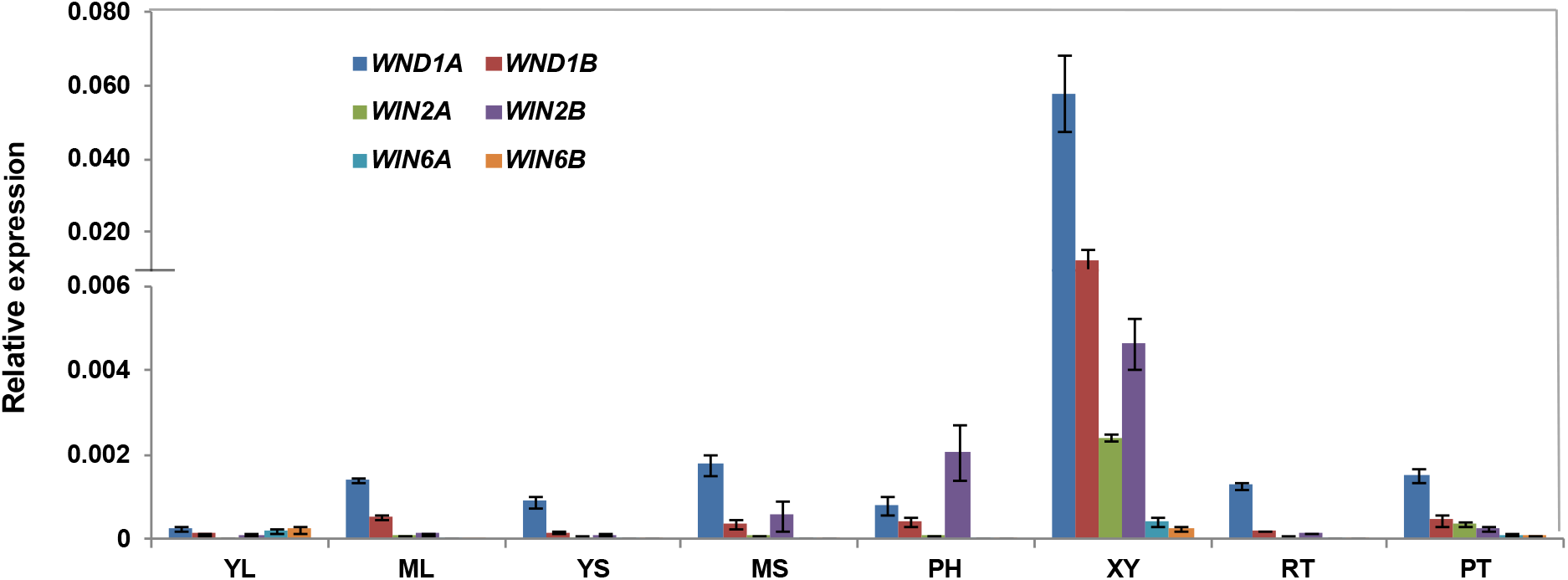
Expression of the six NAC genes in different tissues of *Populus*. YL (young leaf), ML (mature leaf), YS (young stem), MS (mature stem), PH (phloem), XY (xylem), RT (root), PT (petiole). Relative expression (arbitrary units) was calculated based on the expression of target genes relative to house-keeping genes, *Ubiquitin conjugating enzyme E2* and *18S RNA*. A break in the Y-axis represents discontinuous scale. The data represents mean values of three biological replicates ± SE.

**Supplemental File 4.**
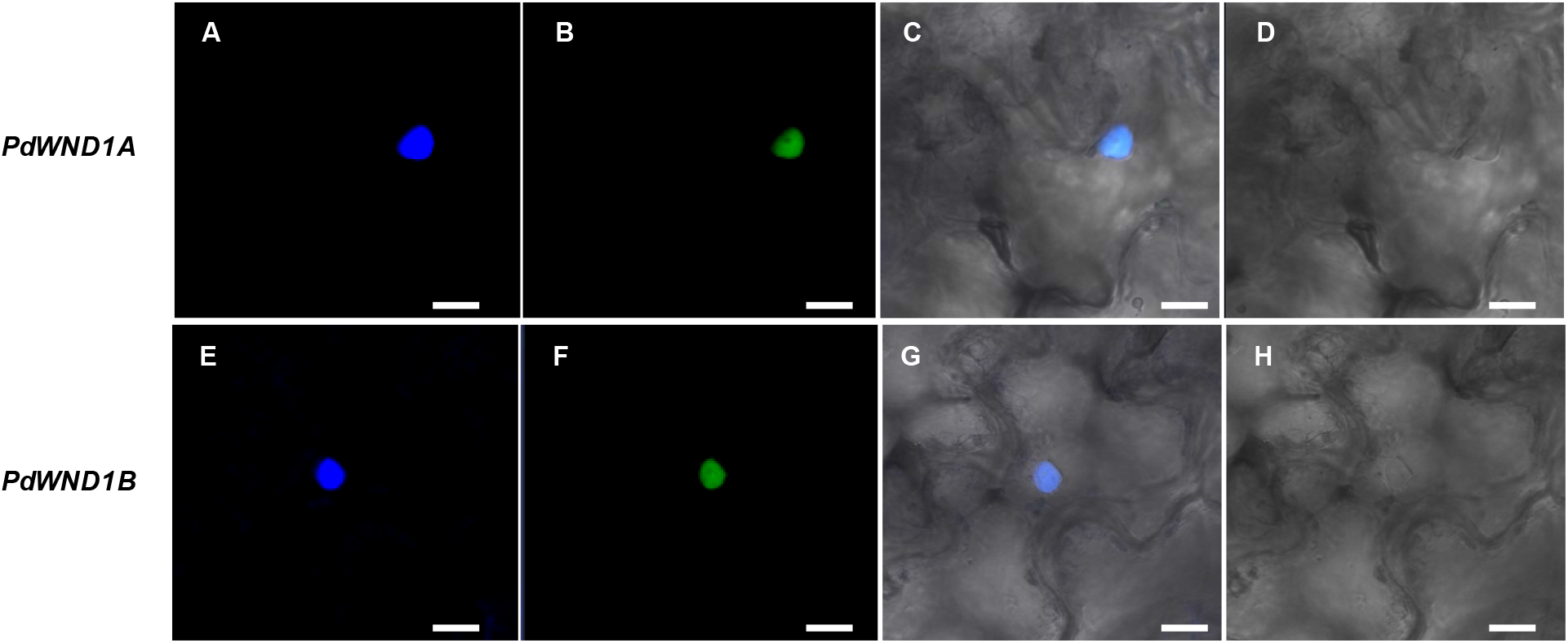
Localization of the PdWND1A and PdWND1B in tobacco epidermal cells. Nuclear targeting of GFP: **PdWND1A** (A to D) and **PdWND1B** (E to H) in *Nicotiana benthamiana* mesophyll cells after agro infiltration. Panels A and E are cells stained with DAPI to show nuclei (blue stain), B and F are GFP localization, C and G are colocalization of DAPI and GFP, and D and H, no florescence control. Scale bar represents 10 μM.

**Supplemental File 5.**
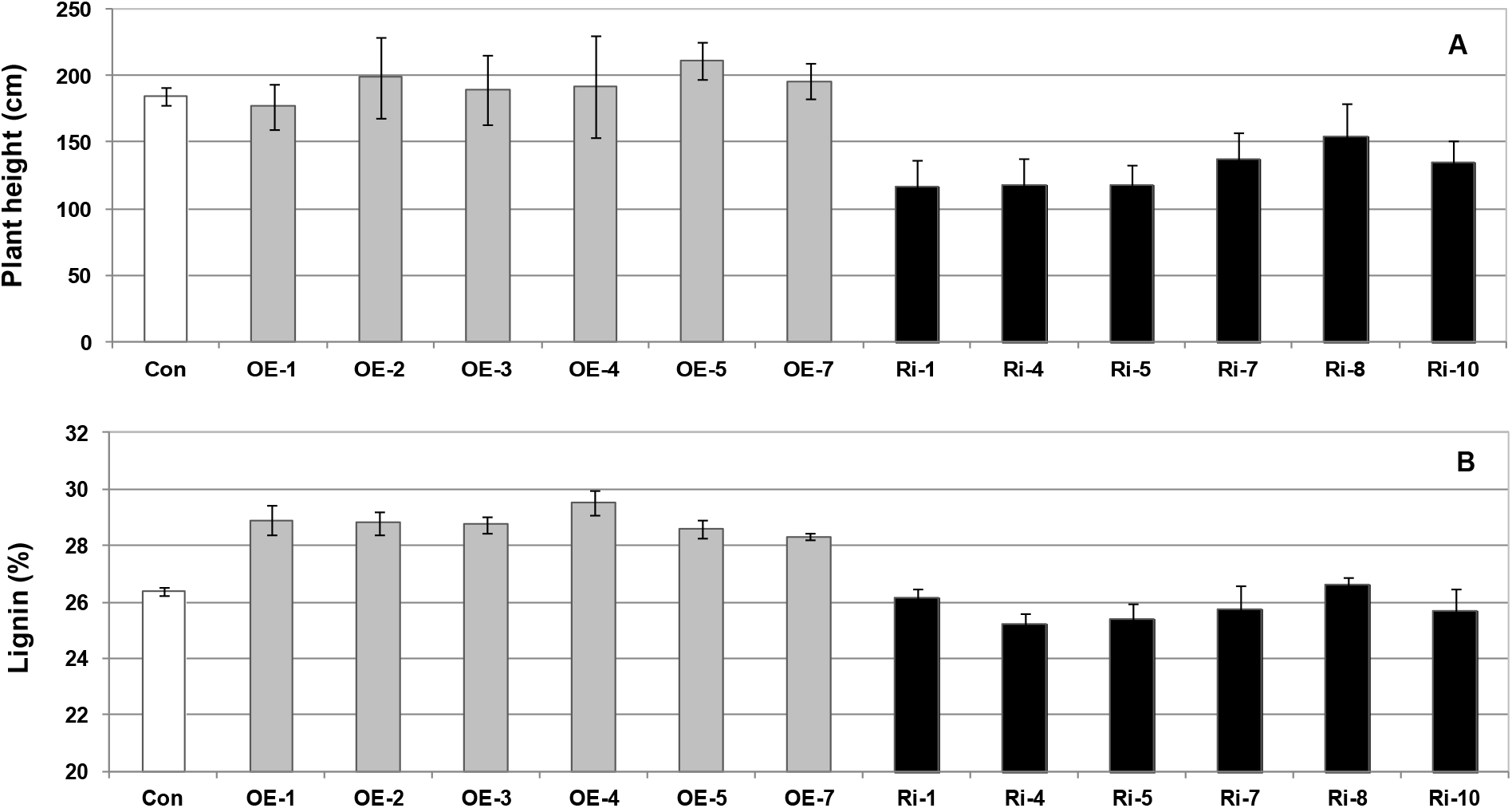
Plant height (A) and lignin content (B) in control (Con) and *PdWND1B* over-expression (OE) and RNAi suppression (Ri) lines.

**Supplemental File 6.**
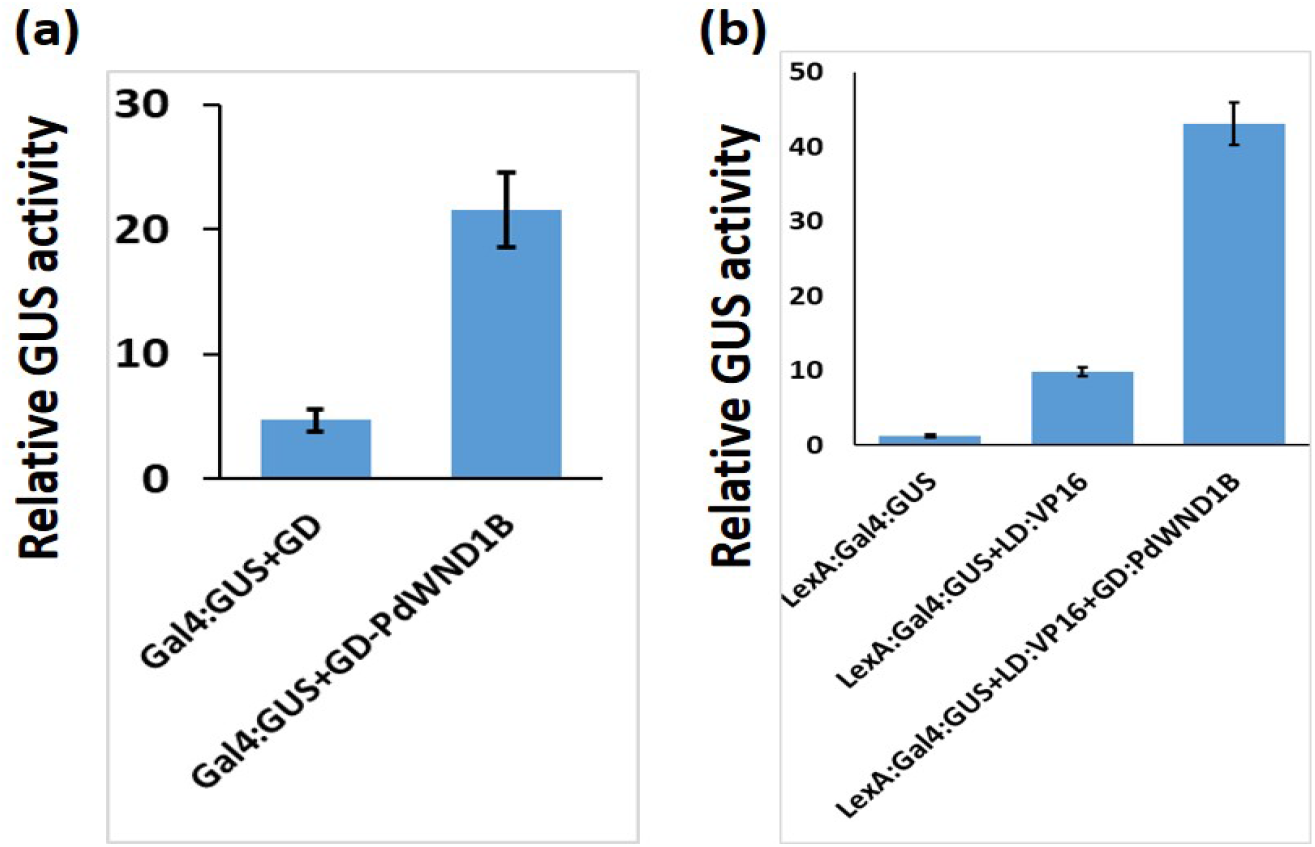
PdWND1B has transcriptional activator activity. (a) Protoplasts transfected with Gal4:GUS reporter together with Gal4 binding domain (GD) fused with PdWND1B (GD-PdWND1B) shows increased GUS activity as compared to empty GD vector control. (b) GD-PdWND1B does not repress the expression of GUS reporter when co-transfected with LexA:Gal4:GUS reporter and LexA binding domain (LD) fused transactivator VP16.

**Supplemental File 7.**
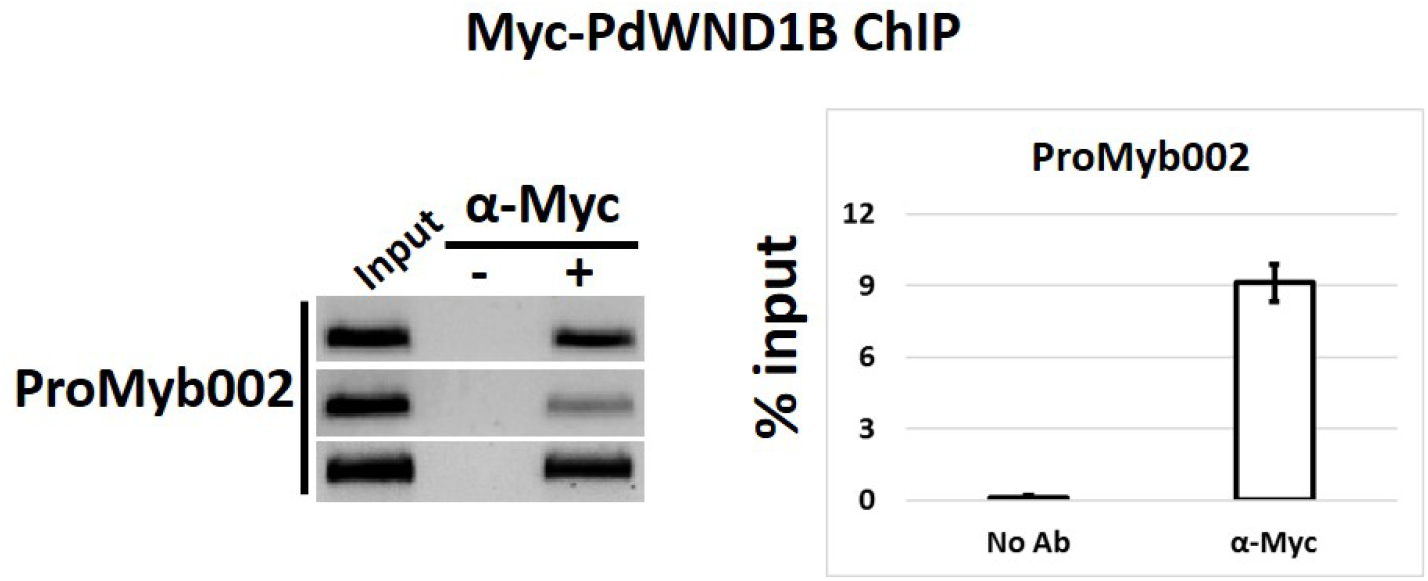
PdWND1B binds to promoter of *MYB002* secondary cell wall transcription factor gene. Miro Chromatin immunoprecipitation (μChIP) from protoplasts transfected with Myc-fused PdWND1B indicates its binding to the promoter region of Myb002 gene in vivo. Left panel shows gel bands from three replicates including the input lane, no-antibody negative control and the sample with antibody. ChIP enrichment signal was calculated from quantitative PCR data as percent of input signal.

**Supplemental File 8.**
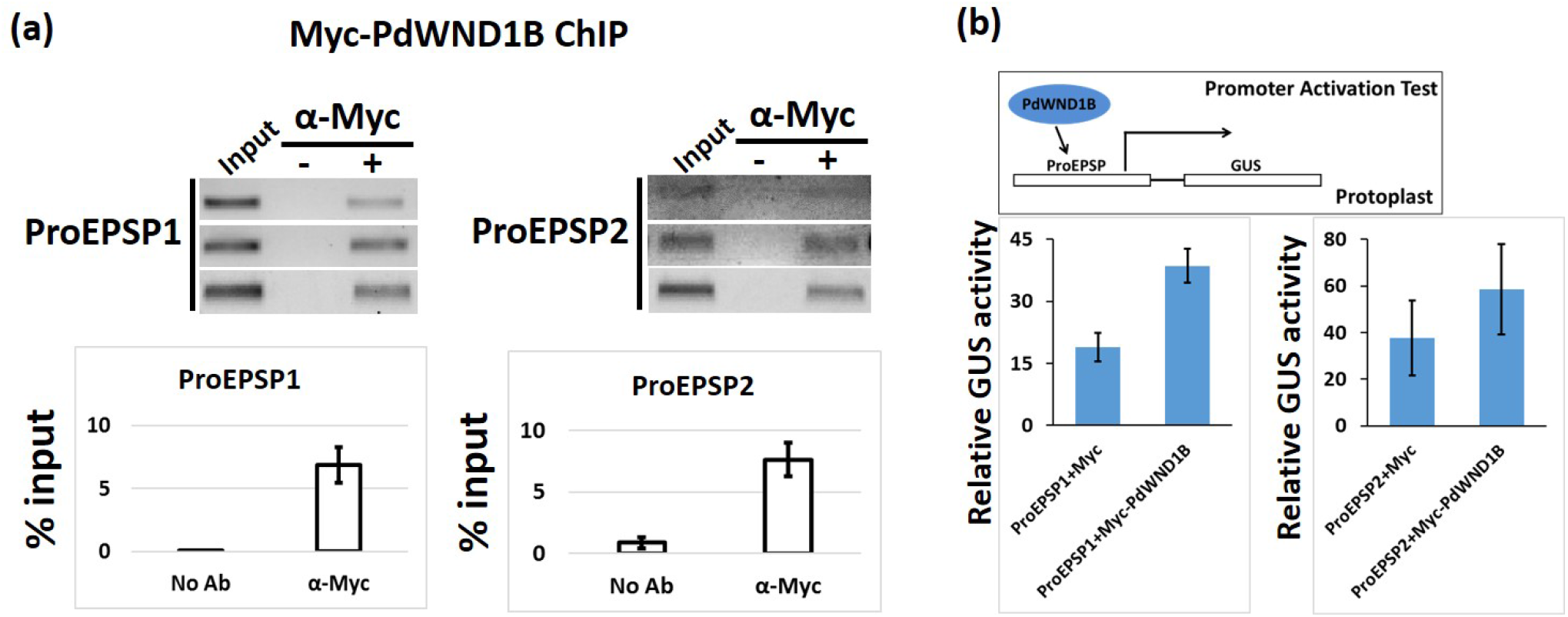
PdWND1B regulates the expression of *EPSP* genes *in vivo*. (a) μChIP from protoplasts transfected with Myc-fused PdWND1B indicates its binding to the promoter region of EPSP1 and EPSP2 genes *in vivo*. Top panel shows gel bands from three replicates including the input lane, no-antibody negative control and the sample with antibody. Bottom panel shows PCR data for ChIP enrichment signal that was calculated as percent of input. (b) Protoplasts transfected with a *EPSP* promoter driven GUS reporter together with the Myc-fused PdWND1B show higher GUS activity as compared to empty vector controls.

**Supplemental File 9.**
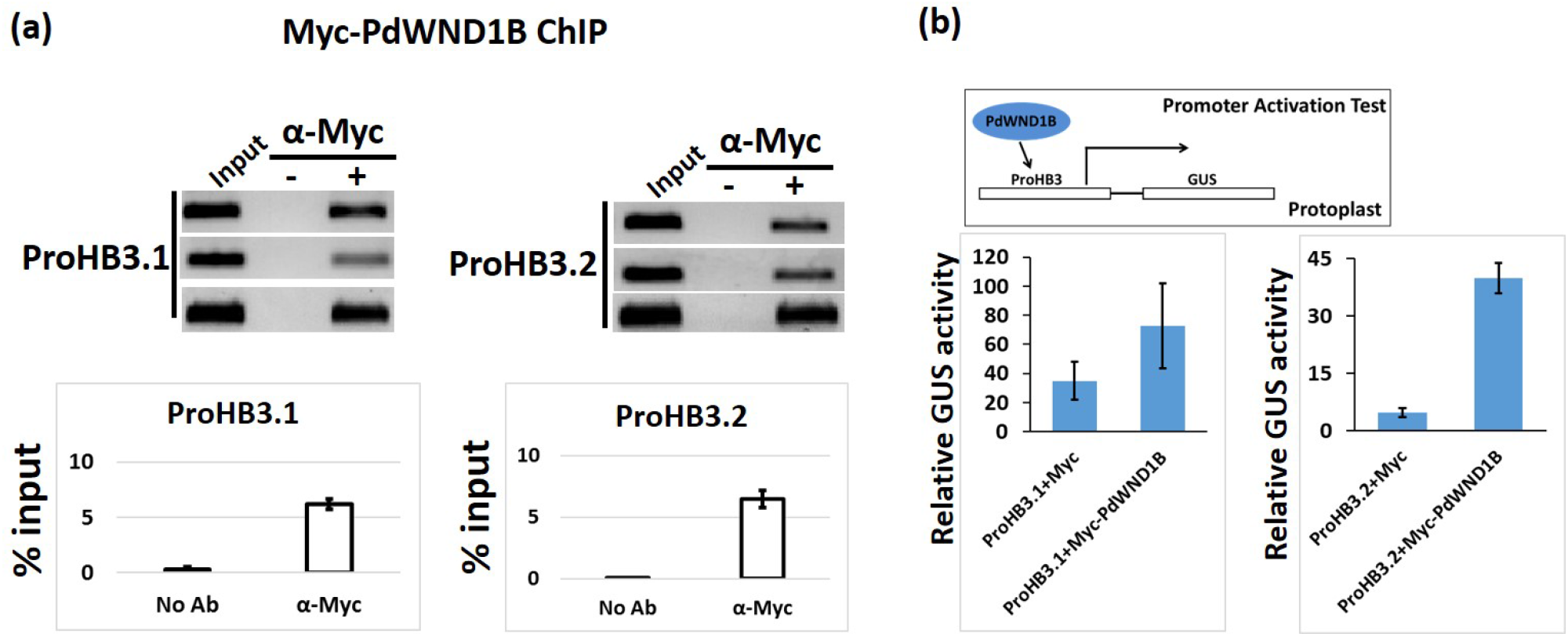
PdWND1B regulates the expression of *HB3-like* genes *in vivo*. (a) μChIP from protoplasts transfected with Myc-fused PdWND1B indicates its binding to the promoter region of *PdHB3-like* genes, *PdHB3.1* (*PtHB3*; Potri.011G098300) and *PdHB3.2* (*PtHB4*, Potri.001G372300), *in vivo*. Top panel shows gel bands from three replicates including the input lane, no-antibody negative control and the sample with antibody. Bottom panel shows PCR data for ChIP enrichment signal that was calculated as percent of input. (b) Protoplasts transfected with an HB3 promoter driven GUS reporter together with the Myc-fused PdWND1B show higher GUS activity as compared to empty vector controls.

